# A conserved pressure-driven mechanism for regulating cytosolic osmolarity

**DOI:** 10.1101/2023.03.01.529730

**Authors:** Katrina B. Velle, Rikki M. Garner, Tatihana K. Beckford, Makaela Weeda, Chunzi Liu, Andrew S. Kennard, Marc Edwards, Lillian K. Fritz-Laylin

## Abstract

Controlling intracellular osmolarity is essential to all cellular life. Cells that live in hypo-osmotic environments like freshwater must constantly battle water influx to avoid swelling until they burst. Many eukaryotic cells use contractile vacuoles to collect excess water from the cytosol and pump it out of the cell. Although contractile vacuoles are essential to many species, including important pathogens, the mechanisms that control their dynamics remain unclear. To identify basic principles governing contractile vacuole function, we here investigate the molecular mechanisms of two species with distinct vacuolar morphologies from different eukaryotic lineages—the discoban *Naegleria gruberi*, and the amoebozoan slime mold *Dictyostelium discoideum*. Using quantitative cell biology we find that, although these species respond differently to osmotic challenges, they both use actin for osmoregulation, as well as vacuolar-type proton pumps for filling contractile vacuoles. We also use analytical modeling to show that cytoplasmic pressure is sufficient to drive water out of contractile vacuoles in these species, similar to findings from the alveolate *Paramecium multimicronucleatum*. Because these three lineages diverged well over a billion years ago, we propose that this represents an ancient eukaryotic mechanism of osmoregulation.

## INTRODUCTION

Osmoregulation is critical for cell survival, and must be tightly controlled. Many species that inhabit hypo-osmotic environments use contractile vacuoles to collect excess water and pump it out of the cell—a similar strategy to bailing water from a leaking boat. Although these organelles are common across eukaryotic phyla (**Fig. 1A**),^1–11^ have been long-studied,^12^ and are considered relevant drug targets in important pathogens,^4, 5, 13^ the mechanisms that drive contractile vacuole pumping remain mysterious.

**Figure 1.**
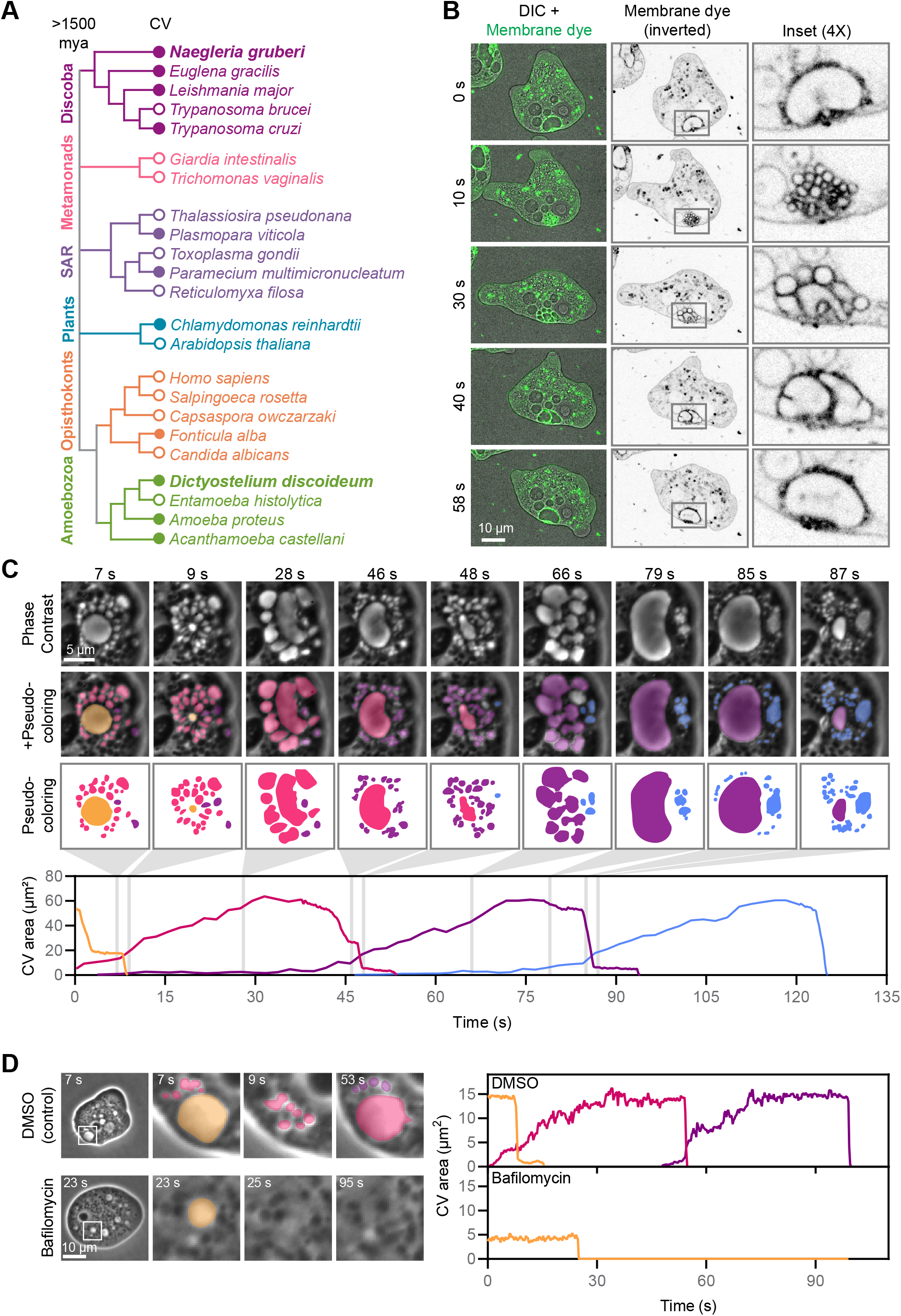
*Naegleria* has a contractile vacuole network that requires vacuolar-type H+ ATPase activity. (**A**) The evolutionary relationships between select eukaryotes are shown using a cladogram, with *Naegleria gruberi* and *Dictyostelium discoideum* in bold. Filled circles indicate species that have been shown to use contractile vacuoles (CVs). (**B**) A representative *N. gruberi* cell stained with the membrane dye FM4-64 highlights the emptying and filling of its contractile vacuole network. (**C**) Contractile vacuole networks were imaged through multiple pump cycles. Vacuoles were tracked as they formed, grew, merged, and collapsed and are pseudocolored by pump (top). The graph shows the area (the 2-D footprint) corresponding to each contractile vacuole network over time. (**D**) Cells were treated with the vacuolar-type H+ ATPase inhibitor Bafilomycin A1 or DMSO (carrier control), and imaged. Contractile vacuole network areas were tracked as in panel C. Also see **Fig. S1D**.

Osmotic regulation by contractile vacuoles can be divided into two major steps: collection of water into a contractile vacuole network, and expulsion of water from the cell. The collection step relies on active transport of solutes and passive flow of water: vacuolar-type H^+^-ATPases (vacuolar-type proton pumps) embedded in contractile vacuole membranes raise the osmotic strength of the lumen, facilitating passive water flow from the cytosol through aquaporins.^3, 14–22^ Contractile vacuole network morphology is dynamic and varies among species. *Paramecium*, for example, has two star-shaped contractile vacuole networks, with “canals” that extend outward into the cytosol to collect water that then flows into a central “bladder.”^23, 24^ In contrast, the contractile vacuole networks of *Dictyostelium* are less uniform, comprising tubules that collect water and expand to form bladders.^9, 16, 25^ Meanwhile, *Amoeba proteus* collects water into clusters of vacuoles that fuse to form bladders.^26^ In all of these networks, the bladders periodically expel their contents into the environment. During expulsion events, the contractile vacuole is maintained as a distinct compartment, which collapses into a membranous network when emptied.^9, 25, 26^ *Paramecium* contractile vacuoles expel water through a stable, dedicated pore,^27^ while *Dictyostelium* forms a transient pore encircled by an actin mesh that prevents intermixing of contractile vacuole and plasma membranes.^28^

The forces that power bladder emptying have been the focus of much speculation. One hypothesis suggests that contractile vacuole membranes prefer to form thin tubules rather than spherical bladders, and this tendency to tubulate pushes water out of the cell.^16, 28–30^ This hypothesis has not been directly tested, although water expulsion is slower in *Dictyostelium* cells lacking proteins that reinforce membrane curvature, consistent with an important role for tubulation in contractile vacuole function.^31^ A second hypothesis, based largely on localization data from *Acanthamoba,^32–35^* invokes the existence of actomyosin networks that surround the bladder and contract to squeeze water out of the cell.^10, 36^ A third hypothesis, supported by analytical modeling of *Paramecium* contractile vacuoles,^6^ suggests that cytoplasmic pressure alone is sufficient to drive expulsion.^11^ To test these hypotheses and identify basic principles underlying contractile vacuole function, we study species from two distinct eukaryotic lineages: the well-studied amoebozoan slime mold *Dictyostelium* and *Naegleria*—a freshwater amoeba that diverged from amoebozoans over a billion years ago^37–39^ (**Fig. 1A**).

In addition to occupying an evolutionary position useful for identifying conserved contractile vacuole mechanisms, *Naegleria* amoebae also have simplified cell biology and potential relevance to human health. *Naegleria* amoebae lack cytoplasmic microtubules,^40–45^ which guide contractile vacuole membrane trafficking in *Dictyostelium,^46^* and anchor the contractile vacuole in a stable location in other species (e.g. *Paramecium^27^*). Contractile vacuoles are obvious in *Naegleria;* even early cell biologists observed 1-3 large contractile vacuole bladders forming from the coalescence of smaller vacuoles.^1, 47^ Studying *Naegleria* contractile vacuoles also has practical value; one *Naegleria* species, the “brain-eating amoeba” *N. fowleri*, causes a deadly brain infection for which we currently lack effective therapeutics.^48^ During infection, *Naegleria* transition from freshwater to cerebrospinal fluid, which is two orders of magnitude higher in osmolarity,^49^ making the contractile vacuole a potential *Naegleria* drug target.

Here, we identify core mechanisms used by both *Naegleria* and *Dictyostelium* to drive contractile vacuole function. We show that *Naegleria*, like *Dictyostelium*, rely on vacuolar-type proton pumps for filling contractile vacuoles. We also show that, although *Naegleria* and *Dictyostelium* respond differently to osmotic challenges, they both use actin networks for osmoregulation, but not for the water expulsion step. Finally, we use biophysical modeling to show that cytoplasmic pressure is sufficient to drive water expulsion in both *Naegleria* and *Dictyostelium*. These findings suggest the potential for a universal mechanism of contractile vacuole pumping.

## RESULTS

### *Naegleria* use asynchronously filling contractile vacuole networks that require vacuolar-type proton pump activity

Unlike *Dictyostelium*,^9, 15, 16, 25, 28, 29, 31, 46, 50–59^ there is limited information available about *Naegleria* contractile vacuoles.^1, 47, 60^ To determine how similar *Naegleria* contractile vacuoles are to those of *Dictyostelium*, we quantified their dynamics using live cell imaging. To this end, we treated *Naegleria gruberi* cells with FM4-64—a membrane dye used in other species to label contractile vacuole membranes^16, 26^—and restricted the contractile vacuole networks to a single focal plane using an agarose overlay (**Fig. 1B**). Consistent with previous descriptions,^1, 47, 60^ we observed membrane-enclosed vacuoles that grew and shrank at regular intervals, with large bladders forming from the merging of smaller vacuoles (**Fig. 1B, Video S1A**). Also in line with prior reports,^1, 47, 60^ these vacuoles were localized to the back of migrating cells (**Fig. S1A**). To track the fate of each compartment of a contractile vacuole network, we imaged cells through multiple pump cycles using phase contrast microscopy at a higher frame rate of 4 frames/sec (**Fig. 1C, Video S1B**). This analysis revealed occasional pauses during the expulsion period, where bladders neither grew nor shrank (*e.g*. **Fig. 1C**, pump 1, 3-8 s). Rather than a single vacuole swelling period followed by a single pumping event, concurrent swelling and shrinking occurred in distinct compartments (**Fig. 1C**, graph); as one large bladder collapses, other, smaller vacuoles grow, and eventually fuse to form the next bladder. This means that multiple generations of contractile vacuoles co-exist, with vacuoles sometimes becoming detectable two pumps in advance of their own pump (**Fig. 1C**, pumps 3 and 4). The identity of the contractile vacuole system appears fluid, with vacuoles occasionally dispersing to occupy distinct spaces in the cell, and dispersed vacuoles sometimes fusing into larger systems (**Video S2**).

We next sought to determine how water moves from the cytosol into contractile vacuole networks. Other species,^14, 19, 20, 22, 61^ including *Dictyostelium*,^15, 16^ transport protons into their contractile vacuoles using vacuolar-type proton pumps, resulting in a contractile vacuole lumen that is hyperosmotic relative to the cytosol.^21^ Aquaporins in the contractile vacuole membrane then allow water to flow passively into the contractile vacuole.^3, 17, 18^ In *Dictyostelium*, vacuolar-type proton pumps are readily visualized using deep etch electron microscopy, where they appear as 10 nm pegs protruding from the contractile vacuole membrane.^16^ Similar pegs have been reported in *Naegleria* as well,^16^ consistent with our identification of a full complement of vacuolar-type H+ATPase subunits along with aquaporins in the *Naegleria* genome (**Table 1, Fig. S1B-C**). To directly test if these proton pumps are required for contractile vacuole function in *Naegleria*, we treated cells with the vacuolar-type H+ATPase inhibitor Bafilomycin A1.^62, 63^ Treatment with 100 nM Bafilomycin A1 prevented the refilling of contractile vacuoles (**Fig. 1D, Fig. S1D**), consistent with vacuolar-type proton pump activity facilitating water flow from the cytosol into the contractile vacuole.

**Table 1.** The *Naegleria* genome contains homologs of aquaporins and a bafilomycin-sensitive v-ATPase. Table of *N. gruberi* genes matching the given PFAM models for aquaporins or subunits of v-ATPase, as well as the top-scoring BLASTP hit to each *N. gruberi* gene in the *S. cerevisiae* or *D. discoideum* genome. Genes are referred to by their NCBI accession ID, as well as the JGI ID (for *N. gruberi*) or a conventional name (for *S. cerevisiae* and *D. discoideum*). Asterisk indicates that V1 subunit G did not return significant matches unless the low complexity filter was removed from the BLASTP settings – see **Fig. S1B** and Methods.

|  | Subunit | Matching PFAM ID(s) | *N. gruberi* gene |  | Top *S. cerevisiae* BLASTP hit |  | Top *D. discoideum* BLASTP hit |  |
| --- | --- | --- | --- | --- | --- | --- | --- | --- |
|  |  |  | JGI | NCBI | name | NCBI | name | NCBI |
| v-ATPase V0 | a | PF01496 | 59034 | XP_002673239.1 | Vph1 | NP_014913.3 | vatM | XP_629892.1 |
|  |  |  | 75402 | XP_002669641.1 |  |  |  |  |
|  | c, c' | PF00137 | 82300 | XP_002668836.1 | Vma11 | NP_015090.1 | vatP | XP_644319.1 |
|  |  |  | 4925 | XP_002678119.1 |  |  |  |  |
|  |  |  | 83282 | XP_002671873.1 |  |  |  |  |
|  | c'' |  | 80877 | XP_002673621.1 | Vma16 | NP_011891.1 | DDB_G0274141 | XP_644318.1 |
|  | d | PF01992 | 81090 | XP_002672968.1 | Vma6 | NP_013552.3 | vatD-2 | XP_644445.1 |
| v-ATPase V1 | A | PF00006, PF16886 | 59416 | XP_002671597.1 | Vma1 | NP_010096.1 | vatA | XP_637351.1 |
|  | B | PF00006 | 55664 | XP_002680466.1 | Vma2 | NP_009685.3 | vatB | XP_642608.1 |
|  | C | PF03223 | 77339 | XP_002678623.1 | Vma5 | NP_012843.1 | vatC | XP_638562.1 |
|  | D | PF01813 | 49328 | XP_002676793.1 | Vma8 | NP_010863.1 | atp6v1d | XP_643919.1 |
|  |  |  | 74068 | XP_002670923.1 |  |  |  |  |
|  |  |  | 75597 | XP_002669461.1 |  |  |  |  |
|  | E | PF01991 | 71627 | XP_002673226.1 | Vma4 | NP_014977.3 | vatE | XP_643473.1 |
|  | F | PF01990 | 32528 | XP_002678706.1 | Vma7 | NP_011534.1 | vatF | XP_645434.1 |
|  | G | PF03179 | 79649 | XP_002677534.1 | Vma10* | NP_011905.1* | vatG* | XP_642039.2* |
|  | H | PF03224, PF11698 | 78643 | XP_002680972.1 | Vma13 | NP_015361.1 | vatH | XP_644034.1 |
| Aqua-porins |  | PF00230 | 78300 | XP_002681822.1 | Aqy3 | NP_116601.1 | aqpB | XP_641629.1 |
|  |  |  | 75649 | XP_002669389.1 | Aqy3 | NP_116601.1 | aqpB | XP_641629.1 |

Because the contractile vacuole membrane and associated vacuolar-type proton pumps facilitate water flow into the vacuole lumen, its surface area likely plays a key role in osmoregulation. Having multiple smaller vacuoles provides a larger surface area than one single compartment of equal volume. To explore whether this increased surface area helps in building and/or maintaining an osmotic gradient, we developed a mathematical model based on mass conservation to derive a relation between the osmotic gradient across the contractile vacuole membrane and the osmotic gradient across the plasma membrane (**Text S1**). This model suggests that an increased surface area allows water to flow into the contractile vacuole at a lower osmotic pressure difference between the cytosol and vacuole lumen. This means that a larger contractile vacuole membrane requires less vacuolar-type proton pump activity— and therefore less ATP—to regulate intracellular osmolarity. We also performed an orders-of-magnitude estimation of the “critical size” at which a contractile vacuole will pump by balancing the energetic gain from lowering the osmotic pressure and the energetic penalty from increasing membrane surface area. Entering reasonable values for membrane tension and cytosolic elastic energy^64–66^ into this equation results in a critical radius of 1 μm, which is consistent with our experimental observations. Taken together, these analyses indicate that the filling of *Naegleria* contractile vacuoles can be explained by a simple energetic balance between the osmotic pressure and membrane tension.

### The contractile vacuoles of *Naegleria* respond differently to environmental changes than those of *Dictyostelium*

Cells typically modulate their contractile vacuole dynamics based on environmental conditions.^7, 59, 67–70^ For example, *Dictyostelium* pumps more water through its contractile vacuoles when in more dilute solutions.^53^ To determine how *Naegleria* contractile vacuole dynamics change under varying environmental conditions, we imaged *Naegleria* at two temperatures (20 °C and 35 °C) and two osmolarities (0 mOsm and 50 mOsm) using sorbitol—a non-metabolizable sugar used to increase osmolarity^6, 59^ (Note: the concentration of sorbitol in mM matches its osmolarity in mOsm). We then quantified the frequency of expulsion events from the largest recurring bladder (pumping rate of main contractile vacuole) and the largest bladder size (maximum CV area) for each of ten cells in five independent experiments (**Fig. 2A**). The pumping rate more than doubled at the higher temperature, regardless of external osmolarity: at 20 °C, contractile vacuoles pumped at an average of 0.9 pumps/min in both 0 and 50 mM sorbitol, while at 35 °C, contractile vacuoles pumped at 2.0 pumps/min in both osmolarities (p=1.1E-9 at 0 mM, p=2.2E-9 at 50 mM). These rates are within the previously reported range of 0.3-2.4 pumps per minute.^1, 60^ Although pumping frequency did not vary with osmolarity, the maximum size of the bladder did: contractile vacuoles were over 3 times larger in 0 mM sorbitol than in 50 mM sorbitol (45.9 vs. 14.5 μm^2^ at 20 °C, p=4.7E-8 and 61.8 vs. 17.8 μm^2^ at 35 °C, p=3E-10). At 35 °C, the vacuoles in 0 (but not 50) mM sorbitol were significantly larger than at 20 °C (p=2.3E-4). To determine the extent to which *Naegleria* can respond to external osmolarity by altering contractile vacuole size but not pumping rate, we repeated these experiments under an extended range of osmolarities, from 0 to 100 mM sorbitol at room temperature (**Fig. 2B**). We found a similar result: the maximum size decreased from 54.5 μm^2^ in 0 mM sorbitol to 7.7 μm^2^ in 100 mM sorbitol (p=2.1E-6), while the pumping rate did not vary significantly (a high of 1.2 pumps/min was measured in 25 mM sorbitol, and a low of 0.9 pumps/min was measured in 100 mM sorbitol). Together, these data indicate that, while *Naegleria* contractile vacuoles respond to temperature primarily by modulating pumping rate, they respond to osmolarity by modulating bladder size.

**Figure 2.**
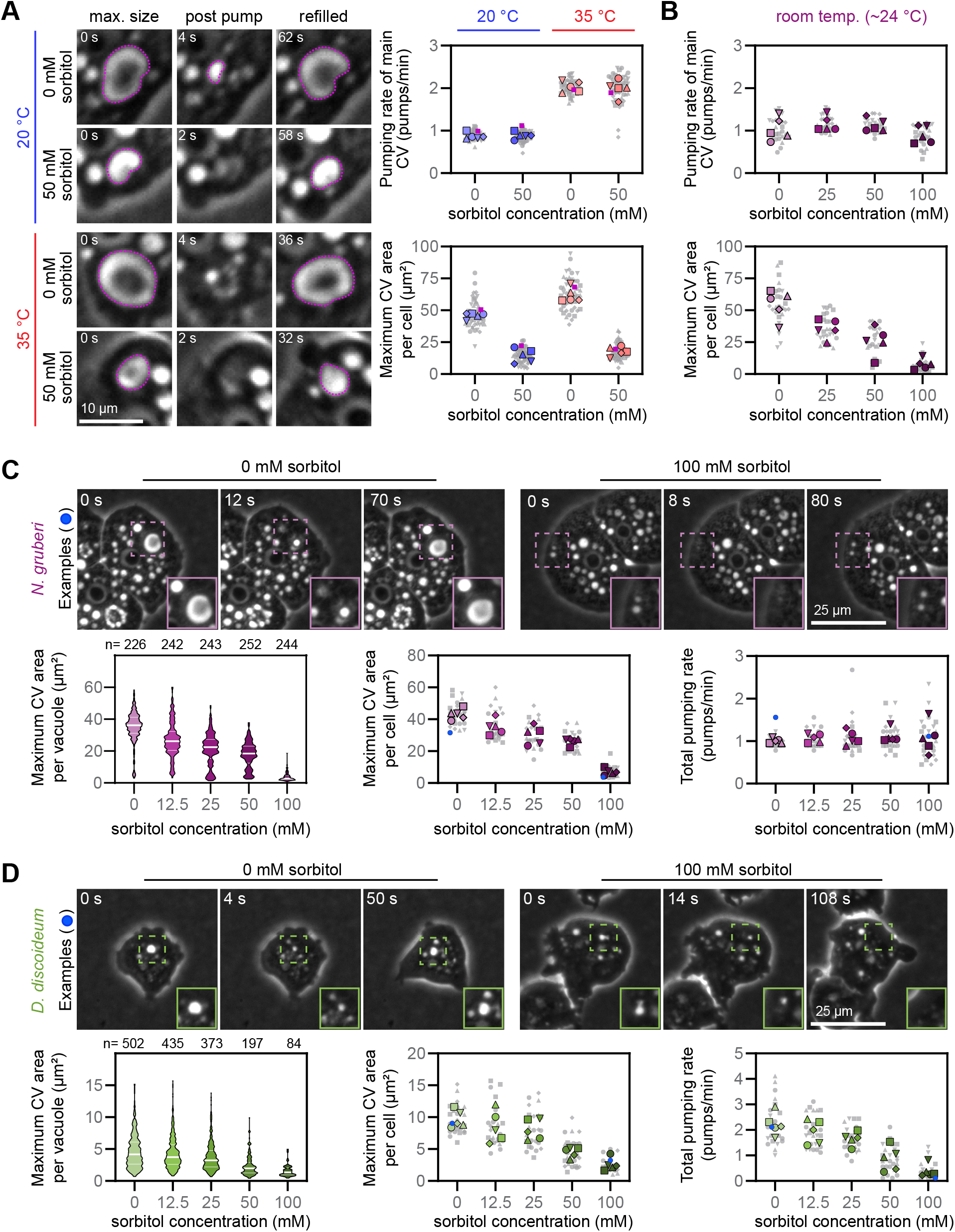
The contractile vacuoles of *Naegleria* respond differently to environmental changes than those of *Dictyostelium*. (**A**) Cells were incubated in water or 50 mM sorbitol, and imaged at low (20+/-1 °C) or high (35+/-1 °C) temperatures. The images show one example cell for each condition with one contractile vacuole bladder outlined in purple for each cell. Contractile vacuoles were quantified to determine the pumping rate of the main bladder (top), as well as the largest contractile vacuole bladder per cell (bottom). Each small gray symbol represents a single cell (10 cells per experiment), while larger symbols represent experiment-level averages for 5 replicates, with symbols coordinated by experiment, and the representative cells in purple. (**B**) Cells were treated with water or the indicated concentrations of sorbitol and imaged as in A, but at room temperature. The pumping rate (top) and maximum contractile vacuole area (bottom) were quantified as in A, with 5 cells per experimental replicate. (**C**) *N. gruberi* cells were exposed to water or the indicated concentration of sorbitol, and imaged. Top panels show example cells with insets magnified at 1.5X. Bottom graphs show the maximum contractile vacuole area for each pumping event (left, violin plot) or cell (center, SuperPlot), and the number of pumping events per minute (right, SuperPlot). The violin plot shows the pooled values for 5 experimental replicates, with white lines indicating the median and quartiles (also see **Fig. S2**). In the SuperPlots, each small gray symbol represents a single cell (5 cells per experiment), while larger symbols represent experiment-level averages for 5 replicates, with symbols coordinated by experiment. The two example cells are highlighted in blue. (**D**) *D. discoideum* cells were exposed to water or the indicated concentration of sorbitol, and imaged and quantified as in A.

Our finding that *Naegleria* maintains a constant contractile vacuole pumping rate under different osmolarities contrasts with *Dictyostelium discoideum*, which has been reported to increase pumping activity as a response to hypo-osmotic stress.^53^ To confirm this difference, we directly compared the contractile vacuole networks of *Naegleria* and *Dictyostelium* in side-by-side experiments. We measured every pumping event in both cell types across a range of sorbitol concentrations (**Fig. 2C-D, Fig. S2**). Both *Naegleria* and *Dictyostelium* cells formed larger vacuoles at lower concentrations of sorbitol. The average maximum vacuole size per cell for *Naegleria* was 43.0 μm^2^ in 0 mM sorbitol and 7.0 μm^2^ in 100 mM sorbitol (p=3.9E-11), while those of *Dictyostelium* ranged from 9.7 μm^2^ in 0 mM sorbitol to only 2.6 μm^2^ in 100 mM sorbitol (p=5.1E-6). Unlike *Naegleria* cells, which maintained a pumping rate of 1.0-1.1 pumps/min across sorbitol concentrations, *Dictyostelium* cells had a higher pumping rate at lower concentrations of sorbitol (2.2 pumps/min at 0 mM, 1.9 at 12.5 mM, 1.7 at 25 mM, 0.9 at 50 mM, and 0.4 at 100 mM sorbitol). Collectively, these data show that while both amoebae adjust vacuole size to account for osmolarity, only *Dictyostelium* pumps more frequently when under higher hypo-osmotic stress.

### Actin is involved in *Naegleria* osmoregulation, but is not required for contractile vacuole pumping

Because actin networks have been hypothesized to drive contractile vacuole pumping in *Acanthamoeba*,^10^ we searched for actin polymers at *Naegleria* contractile vacuoles in samples subjected to quick-freeze deep-etch electron microscopy. Although we did observe actin filaments at the cell cortex, we did not observe any actin filaments at contractile vacuole membranes (**Fig. 3A**), a similar result to what has been described for *Dictyostelium.^16^* Although the lack of actin at contractile vacuoles strongly suggests that actin network contraction cannot drive water expulsion from contractile vacuoles, it is possible that actin assembly is transient and difficult to detect using electron microscopy. Therefore, we treated cells with 5 μM latrunculin B (LatB)—which results in global actin depolymerization^71, 72^ (**Fig. S3A**)—and measured contractile vacuole activity. Strikingly, this treatment did not eliminate contractile vacuole pumping (**Fig. 3B, Video S3**). The ability of these vacuoles to continue pumping in the absence of actin filaments is incompatible with the actomyosin contractility hypothesis that relies on the existence of an actin network.

**Figure 3.**
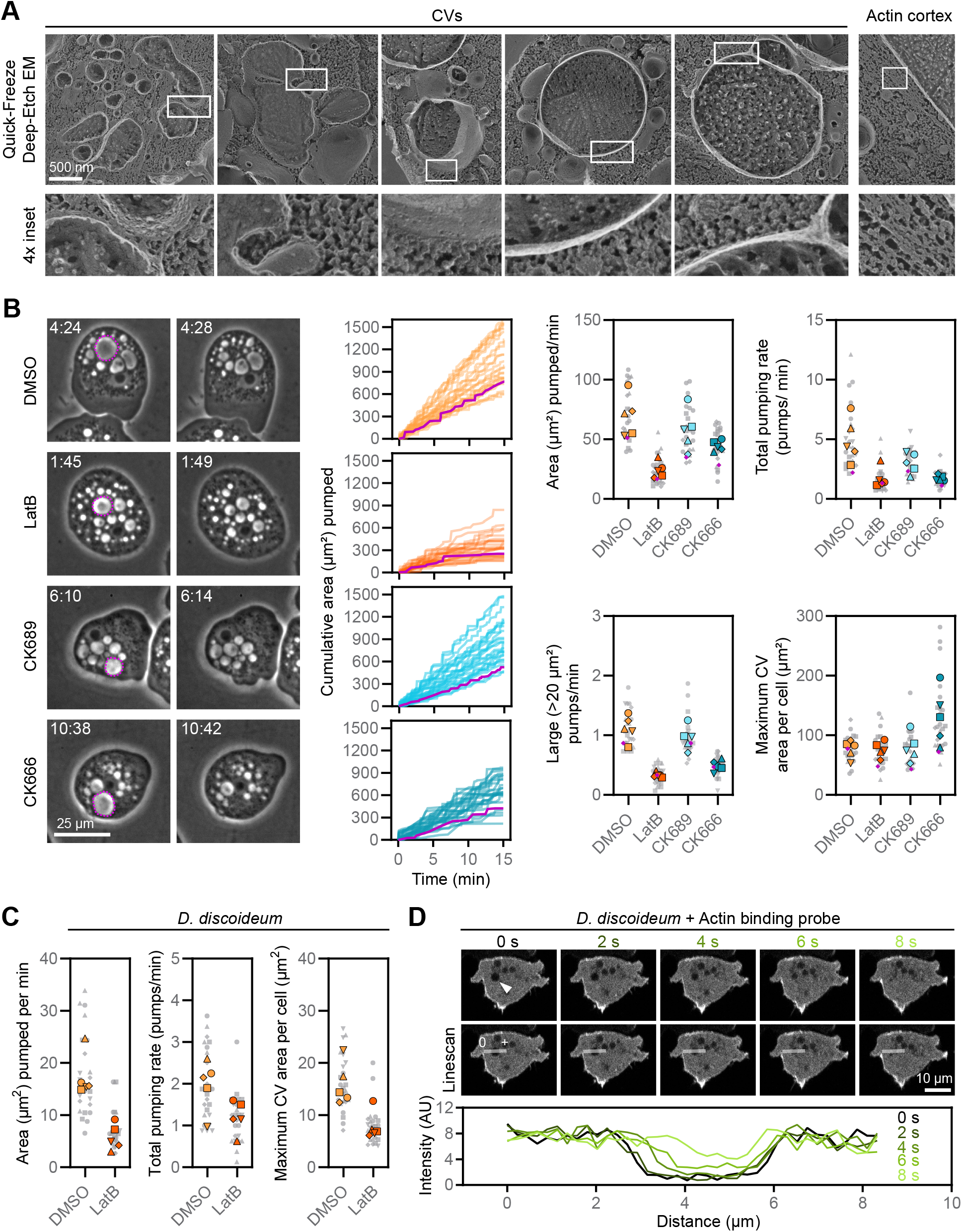
Actin is involved in osmoregulation but is not required for contractile vacuole pumping events. (**A**) *N. gruberi* cells (strain NEG) were subjected to Quick-Freeze Deep-Etch Electron Microscopy. Representative contractile vacuoles and a section of the actin cortex are shown. (**B**) *N. gruberi* cells were incubated in water with the indicated small molecules and controls for ~20 min, then imaged for 15 min. Each contractile vacuole bladder was measured at its maximum size, and the cumulative contractile vacuole area was calculated over the imaging time. Left panels show examples of pumping events when the largest contractile vacuole was present (dashed outline; time is in min:s after imaging began). Middle panels show the cumulative area pumped out of the cell for 25 cells: 5 cells each from 5 experiments, with example cells from the left highlighted in magenta. Right panels show the area pumped per minute, the total number of pumping events per minute, the number of pumping events above a threshold of 20 μm^2^ per minute, and the maximum contractile vacuole area for each cell. Each small gray symbol represents a single cell (5 cells per experiment), while larger symbols represent experiment-level averages for 5 replicates, with symbols coordinated by experiment. The example cells are highlighted in magenta. (**C**) *D. discoideum* cells were incubated in water and treated with LatB or DMSO (vehicle control), and imaged and quantified as in A. The average area pumped per minute, the total number of pumping events per minute, and the maximum contractile vacuole area for each cell are shown. See **Fig. S3B** for additional inhibitors. (**D**) A *D. discoideum* cell expressing a fluorescently-labeled actin-binding protein (RFP-LimE) is shown through a pumping cycle. Line scans bisecting a contractile vacuole show no enrichment of actin around the pumping bladder. See **Fig. S3C** for an additional example.

Although *Naegleria* contractile vacuoles clearly do not require actin to pump, cells treated with actin inhibitors do have defects in osmoregulation. To quantify these defects, we estimated the volume of water expelled over time by summing the maximum bladder sizes from each pumping event (**Fig. 3B**). Treating cells with LatB resulted in a ~65% reduction in water pumped compared to a DMSO carrier control (as measured by contractile vacuole area pumped over time: 24.4 μm^2^/min vs. 69.8 μm^2^/min, p=2.0E-4). To explore the nature of this defect, we treated cells with 50 μM CK-666, which inhibits the Arp2/3 complex to specifically block branched actin network assembly.^73, 74^ This resulted in ~23% drop in water pumped out that was not statistically significant compared to the inactive control CK-689 (44.5 μm^2^/min vs. 57.8 μm^2^). These differences must be due to changes in contractile vacuole numbers, sizes, and/or pumping rates. We therefore quantified each parameter and found that compared to DMSO-treated cells, LatB-treated cells have fewer pumping events per minute (5.0 pumps/min for DMSO and 1.7 pumps/min for LatB, p=1.6E-3)—a difference that is exacerbated when looking only at large (>20 μm^2^) pumping events, which were relatively rare in LatB-treated cells (1.1 pumps/min for DMSO and 0.3 pumps/min for LatB, p=3.1E-6). Meanwhile, although CK-666-treated cells had fewer large pumping events than control cells (0.5 pumps/min for CK-666 vs. 1.0 pumps/min for CK-689, p=1.1E-3), these included some of the largest pumping events observed in this study, with some topping 200 μm^2^. These results show that, although not required for contractile vacuole pumping, actin is important for osmoregulation and the reformation of large vacuoles. This may be due to a requirement for cortical actin to prevent loss of contractile vacuole membranes to the plasma membrane, similar to what has been described for *Dictyostelium*.^28^

### Actin networks are also dispensable for *Dictyostelium* contractile vacuole emptying

The lack of visible actin polymers around *Dictyostelium* contractile vacuoles in deep-etch electron micrographs^16^ suggests that *Dictyostelium* contractile vacuoles may also empty using an actomyosin-independent mechanism. We therefore directly tested the requirement for actin polymers in *Dictyostelium* contractile vacuole pumping by treating cells with LatB. Similar to *Naegleria*, depolymerizing *Dictyostelium* actin^75^ did not eliminate contractile vacuole pumping (**Fig. 3C, Fig. S3B**). As a final test of the actomyosin contractility hypothesis, we imaged cells transfected with RFP-LimE, a probe for polymerized actin. While contractile vacuoles were readily observed as voids that routinely grew and shrank, there was no detectable enrichment of actin at these sites as vacuoles filled or emptied (**Fig. 3D, Fig S3C, Video S4**). The Latrunculin data and this live imaging support an actin-independent mechanism for contractile vacuole emptying.

Although actin is not required for expulsion events, the cytoskeleton of *Dictyostelium* has been implicated in osmoregulation. *Dictyostelium* contractile vacuoles use microtubules to traffic through the cell, and a type V myosin to engage the actin cortex.^46^ We therefore directly tested the role of cytoskeletal polymers in *Dictyostelium* osmoregulation by treating cells with a panel of actin and microtubule inhibitors (**Fig. S3B**). Consistent with a role for the cytoskeleton in contractile vacuole membrane trafficking, LatB-treated cells expelled ~67% less water compared to controls, while CK-666-treated cells pumped out ~45% less, and nocodazole-treated cells pumped out ~36% less water (DMSO: 17.4 μm^2^/min; LatB: 5.7 μm^2^/min, p=1.0E-4; CK-666: 9.6 μm^2^/min, p=5.2E-3; nocodazole: 11.1 μm^2^/min, p=2.6E-2). Taken together, these data show that, like *Naegleria*, *Dictyostelium* uses actin for osmoregulation but not to drive water expulsion from the contractile vacuole.

### Biophysical modeling suggests that cytoplasmic pressure is sufficient to power water expulsion from *Naegleria* and *Dictyostelium* contractile vacuoles

Having ruled out actomyosin-mediated contraction, we next explored other mechanisms that could explain the rapid emptying of *Naegleria* and *Dictyostelium* contractile vacuoles. Physical models of fluid expulsion from *Paramecium multimicronucleatum* contractile vacuoles have suggested that cytoplasmic pressure alone can provide sufficient force to empty the contents of a vacuole within seconds.^6^ We therefore wondered whether cytoplasmic pressure could also be driving vacuole emptying in *Naegleria* and/or *Dictyostelium*. To test this possibility, we developed a model (**Fig. 4A**) based on the previous *Paramecium* work.^6^ Here, we define cytoplasmic pressure as the pressure on the cell periphery due to high intracellular concentrations of ions, osmolytes, and macromolecules. This pressure is a combination of both osmotic and oncotic (crowding) pressure.^76^ Under this model, we assume all of the contents of the vacuole exit through a pore of defined size (**Fig. S4**). Flow through this pore is driven by the pressure difference on either side of the pore, meaning that increasing cytoplasmic pressure increases the emptying rate. The pore itself resists flow, such that having a longer, thinner pore reduces the emptying rate. Using this model, we calculate that cytoplasmic pressure can expel the entire contents of a contractile vacuole within one second for a wide range of biologically-relevant cytoplasmic pressures and pore sizes, a finding that is consistent with our measurements of *Naegleria* contractile vacuole emptying rates (**Fig. 1C-D, Fig. S4**). This model also predicts that the rate of vacuole expulsion is constant in time, meaning the cross-sectional area decreases linearly (**Fig. 4A**). To test this prediction, we measured *Naegleria* and *Dictyostelium* contractile vacuole areas through expulsion events. In both species, the cross-sectional area decreased linearly with time (**Fig. 4B-C**), consistent with our cytoplasmic pressure model. Taken together, we find that cytoplasmic pressure is a plausible mechanism for the rapid expulsion of water from contractile vacuoles.

**Figure 4.**
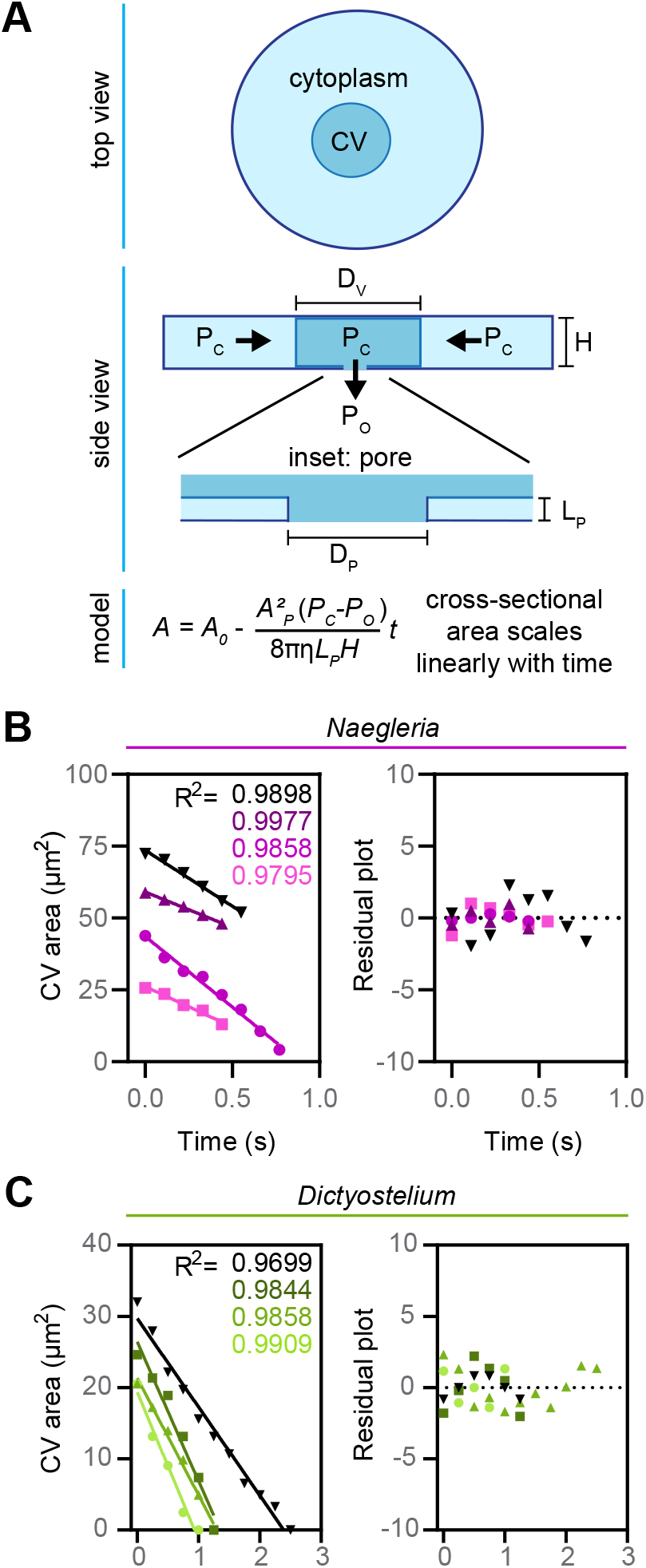
A biophysical model of cytoplasmic pressure-driven contractile vacuole emptying is consistent with contractile vacuole expulsion rates measured in *Naegleria* and *Dictyostelium*. (**A**) Schematic of the model. *Top:* A view of the cell from above. The contractile vacuole (darker circle) is treated as a cylinder sitting inside of a cylindrical cell (lighter circle). *Middle:* A side view of a cell. The contractile vacuole has a diameter, *D_V_*, and a height, *H*, which is equal to the height of the cell. Cytoplasmic pressure, denoted *P_C_*, squeezes water out of the contractile vacuole through a pore. The pressure inside the vacuole is assumed to be equal to the cytoplasmic pressure. As water exits through the pore, the diameter of the contractile vacuole begins to shrink while the height of the contractile vacuole remains constant. *Inset:* A side-view of the pore with diameter *D_P_* and width *L_P_*. *Bottom*: An equation describing the reduction in the contractile vacuole cross-sectional area, *A*, over time, *t*, as water leaves the vacuole through the pore. The contractile vacuole area is described as a function of the initial vacuole area *A*_0_, the area of the pore *A_P_*, the pressure difference across the pore (*P_C_* - *P_O_*), the viscosity of water *η*, the length of the pore *L_P_*, and the height of the contractile vacuole, *H*. (**B**) *N. gruberi* amoebae were imaged at a rate of 0.11 s/frame, and vacuole area was measured for each frame during expulsion. Periods of rapid expulsion were isolated using a threshold of 14 μm^2^/s, and the first time point when rapid expulsion occurred was set to 0. Vacuoles with expulsions lasting for at least 5 consecutive timepoints were analyzed. On the plot of contractile vacuole area over time (left), each point represents the area of the contractile vacuole bladder (n=4 vacuoles from 4 different cells), and the linear regression trendline and R^2^ values are shown. The residual plot (right) is shown to demonstrate the suitability of the linear model. (**C**) The contractile vacuoles of *D. discoideum* cells were imaged at a rate of 0.25 s/frame and quantified as in B, but with a threshold of 4.9 μm^2^/s.

## DISCUSSION

Osmoregulation is critical for cell survival and must be tightly controlled. Here, we show that *Naegleria* and *Dictyostelium* amoebae osmoregulate using contractile vacuole networks that respond to environmental changes. We also show that, like other species, *Naegleria* use vacuolar-type proton pumps for contractile vacuole filling. We confirm that although actin plays roles in osmoregulation, it is dispensable for expelling water from contractile vacuoles of both *Naegleria* and *Dictyostelium*. Instead, our analyses indicate that cytoplasmic pressure is sufficient for this process, similar to findings from the alveolate *Paramecium multimicronucleatum*.^6^ Together, these data reveal simple, conserved mechanisms that underlie contractile vacuole function in diverse eukaryotic species.

Our detailed quantification of contractile vacuole pumping shows that once *Naegleria* bladders begin to empty, they never again swell, suggesting that water may no longer be flowing into these compartments. This raises the possibility that cytosolic water concentration could spike during pumping events. This possibility is ameliorated by a key feature of *Naegleria’s* contractile vacuole network—a division of labor: when one contractile vacuole bladder is pumping water out of the cell, other vacuoles are filling. This asynchronous activity allows for constant water flow out of the cytosol, a potentially conserved feature. For example, *Chlamydomonas reinhardtii* has two bladders that take turns,^77^ and *Paramecium* has canals that continue to fill while the bladder empties.^24^

Exploring the responses of contractile vacuoles to environmental changes also reveals new directions for future research. Similar to other species,^67–69^ *Naegleria’s* contractile vacuoles pump more frequently at higher temperatures, a finding that is consistent with the hypothesis that the historical increase in reported pumping rates of *Naegleria* contractile vacuoles may be due to improvements in laboratory heating.^78^ Unlike other protists,^7, 53, 68^ however, *Naegleria* maintains a constant contractile vacuole pumping rate in different osmotic environments. In side-by-side comparisons, *Dictyostelium* quintupled its pumping rate in water compared to 100 mM sorbitol, while *Naegleria* continued to pump about once per minute in both solutions. This raises an obvious question: how does *Naegleria* maintain a constant contractile vacuole pumping rate? One possible answer may lie in work from *Paramecium* that showed regular membrane tensing of surgically-removed contractile vacuoles,^79, 80^ hinting at the existence of a membrane-associated cycling activity. At the molecular scale, pump frequency may be regulated by ion channels, which are critical for pacemaking activity in cardiac and neuronal cells.^81–83^

Although not required for pumping, actin networks do contribute to osmoregulation, as the contractile vacuoles of Latrunculin-treated cells expelled 65% less water in *Naegleria* and 67% less in *Dictyostelium* compared to controls. This defect could be due to impaired contractile vacuole membrane trafficking,^46^ potentially combined with differences in actin networks at pore sites. In *Dictyostelium*, the actin mesh encircling the pore acts as a barrier to prevent intermixing of contractile vacuole and plasma membranes.^28^ Without an actin cortex, contractile vacuole membranes may completely fuse with the plasma membrane, thereby reducing the available volume of the contractile vacuole network. On the other hand, having too much actin may also impede osmoregulation. CK-666 treatment—which results in a thicker actin cortex in *Naegleria* cells^72^—could block the contractile vacuole from contacting the plasma membrane. This would explain the slower pumping rate and large contractile vacuole bladders we observed upon treating *Naegleria* with CK-666 (**Fig. 3B**).

While our data confirm previous descriptions of *Naegleria* contractile vacuole dynamics,^1, 47, 60^ they refute previous hypotheses of how contractile vacuoles pump. Despite the popularity of the actomyosin contractility hypothesis in the literature,^10, 13, 32, 34–36, 84^ we and others failed to observe actin around active contractile vacuole bladders in any species,^16, 85–88^ and disassembly of all actin networks in *Naegleria* and *Dictyostelium* failed to halt water expulsion. Furthermore, analytical modeling of this mechanism is unable to explain the dynamics of contractile vacuole emptying in *Paramecium.^6^* These data do, however, fit the cytoplasmic pressure model,^6^ as do our biophysical modeling of expulsion events in *Naegleria* and *Dictyostelium*.

Based on these data, we propose that there are only three requirements for pressure-driven contractile vacuole systems: (1) a membrane-bound vacuole that can connect to the cell exterior, (2) a system for transporting water up a gradient, *e.g*. vacuolar-type proton pumps, and (3) cytoplasmic pressure that can push on the contractile vacuole membrane. The simplicity of this system raises the possibility that contractile vacuole networks may be even more widely distributed amongst eukaryotic phyla than is currently appreciated. We now know that cytoplasmic pressure is sufficient to drive contractile vacuole emptying in at least three eukaryotic lineages that diverged from each other over a billion years ago, suggesting that this may represent an ancestral eukaryotic mechanism for regulating intracellular osmolarity.

## METHODS

### Cell culture

*Naegleria gruberi* cells (strain NEG-M, a gift from Dr. Chandler Fulton) were cultured axenically in M7 Media (10% FBS + 45 mg/L L-methionine + 5 g/L yeast extract + 5.4 g/L glucose + 2% (v/v) M7 Buffer (18.1 g/L KH2PO4 + 25 g/L Na2HPO4)) in plug seal tissue culture-treated flasks (CELLTREAT; cat. no. 229330) and grown at 28°C. Cells were split into fresh media every 2-3 days for a maximum of 30 passages. *Naegleria gruberi* strain NEG was used for Quick-Freeze Deep-Etch Electron microscopy and grown on lawns of *Klebsiella pneumoniae* bacteria as previously described.^78^

*Dictyostelium discoideum* cells (strain Ax2-ME, cloned and expanded from AX2-RRK DBS0235521 obtained from the Dictybase stock center)^89^ were grown axenically in HL5 Media (14g/L peptone + 7g/L yeast extract + 13.5 g/L glucose + 0.5g/L KH2PO4 + 0.5g/L NA2HPO4) supplemented with streptomycin (300 μg/mL) and ampicillin (100 μg/mL) on tissue culture-treated plastic 100 mm or 150 mm plates (Fisherbrand Tissue Culture Dish) at 21°C and maintained at 50% confluency by subculturing. For transformation and exogenous gene expression of the plasmid RFP-LimE (Dictybase.org), cells were electroporated as previously described.^90^ Briefly, cells were washed twice and resuspended in H-50 buffer (16.5mM HEPES, 50mM KCl, 10mM NaCl, 1mM MgSO4, 5mM NaHCO3, and 1mM NaH2PO4, pH 7). 5-6 million cells were transferred to a cold 0.1cm electroporation cuvette containing 2 μg of the plasmid and electroporated twice (0.85kV, 25μF) with a 5 s gap between pulses. Electroporated cells were maintained under selection in HL5 supplemented with 20μg/mL G418.

### FM4-64 membrane staining

To visualize cell membranes (**Fig. 1B**), ~3X10^5^ NEG-M cells were seeded into a 6-well glass-bottom plate (Cellvis, P06-1.5H-N). FM™ 4-64 Dye (Invitrogen, T13320) was added to a final concentration of 50 μg/mL. A 1.5% agarose pad (without dye, see under agarose assays below) was overlaid, and the media/staining solution was aspirated off the cells. Cells were imaged in DIC and fluorescence, with images captured every 1.58 s for several pumping cycles. See the microscopy section for details.

### Quantification of contractile vacuole area over time

~3X10^5^ cells were seeded into a 6-well glass-bottom plate (Cellvis, P06-1.5H-N), rinsed with 1 mL of water, and confined under a 1.5% agarose pad (see under agarose assays below). Images were acquired every 250 ms, for multiple pumping cycles (see microscopy section for details). Pumping events were quantified in Fiji^91^ by outlining contractile vacuole bladders at their largest (using the wand tracing tool, or the freehand selection tool for less clear bladders), then measuring the vacuole size every 250 ms as it shrank until it was undetectable, and also working backward from the maximum size to measure the combined area of every smaller vacuole that fused to form that bladder every 2 s (**Fig. 1C**). This information was used to color code the vacuoles by pump using Adobe Illustrator (**Fig. 1C**) or Adobe Photoshop (**Video S1B**). A similar analysis was performed for **Fig. 1D**, but with measurements every 250 ms.

To quantify contractile vacuole expulsion (**Fig. 4B-C**), cells were treated as above and videos of *N. gruberi* were taken at a rate of 110 ms/frame, and videos of *D. discoideum* were taken at a rate of 250 ms/frame. 10 cells were quantified for each species; the area of the bladder was determined at its largest (using the wand tracing tool or freehand selection tool), and the vacuole area was measured in each frame until it was no longer detectable. Because contractile vacuoles sometimes pause during expulsion, periods of rapid expulsion were isolated for analysis using a threshold of 14 μm^2^/s for *N. gruberi* and 4.9 μm^2^/s for *D. discoideum*. Cells that had rapid expulsion events encompassing at least 5 data points were used to analyze the linearity of contractile vacuole area over time.

### Homolog Identification

To identify *N. gruberi* v-ATPase subunit homologs, PFAM models corresponding to each of the 13 subunits were extracted from the Pfam-A database (v35.0) and used to search a curated list of predicted *N. gruberi* proteins (Naegr1_best_proteins.fasta available at https://genome.jgi.doe.gov/portal/Naegr1/Naegr1.download.ftp.html), using HMMER v3.3.2 with default gathering thresholds to filter matches. A BLASTP search was performed by comparing each of the resulting *N. gruberi* genes to proteins from *Saccharomyces cerevisiae* strain S288C (NCBI taxid: 559292) and *Dictyostelium discoideum* strain AX4 (NCBI taxid: 352472), using a word size of 3 and the BLOSUM45 scoring matrix, with all other settings set to their default. Only the top-scoring hit from each species was reported. The score for all reported hits was greater than 74 and the E-value was less than 1e-17. When running this BLASTP search, no significant hits were found matching the *N. gruberi* subunit G (JGI ID: 79649). This may be due to the small size of the protein and its low sequence complexity. A second BLASTP search was conducted for this protein removing the low complexity filter, which returned the reported hits, which had a score greater than 44 and an E-value less than 3e-7. A reciprocal BLASTP search, with the low-complexity filter applied using the *S. cerevisiae* homolog as bait, returned both *N. gruberi* and *D. discoideum* hits. The sequences were aligned with T-COFFEE using default settings and displayed with JalView, using the Clustal coloring scheme with a conservation threshold of 30. *N. gruberi* aquaporins were identified using HMMER, scanning the curated list of *N. gruberi* proteins for matches to the Major Intrinsic Protein PFAM model (PF00230) that passed default gathering thresholds. Examination of these matches with InterProScan only returned related domain predictions (e.g. transmembrane domains or aquaporin-like domains, IPR023271). A BLASTP search using the same settings described above was performed to return the closest matches in *S. cerevisiae* or *D. discoideum*, which had scores greater than 57 and E-values less than 2e-9.

### Measurement of contractile vacuole orientation

~1X10^5^ *N. gruberi* cells were seeded into a 6-well glass-bottom plate (Cellvis, P06-1.5H-N), rinsed with 2 mL of water, and confined under a 1.5% agarose pad (see details on under agarose assays below). Cells were imaged at a rate of 1 frame/s for 5 min.

A semi-automated procedure was developed to quantify the contractile vacuole orientation across multiple cells over time. Briefly, phase contrast movies of sparsely seeded *N. gruberi* cells under agarose were manually contrast-adjusted, and the initial positions of cells were recorded manually.

Cells were automatically segmented using an edge-detection filter and Otsu segmentation, followed by a watershed algorithm and refinement of the cell boundary using active contours. Cells were tracked in the process of segmentation using the centroids of cells in frame *t* – 1 as seeds for cell segmentation in frame *t*. Once cells were segmented and tracked, the centroid velocity was calculated from the total displacement over 5 frames. Contractile vacuole locations at the moment of pumping were identified manually using Fiji, and each pump was associated with the corresponding cell. The orientation of the contractile vacuole was then defined as the angle between the cell to contractile vacuole centroid-to-centroid vector and the cell centroid velocity vector, with the origin of both vectors at the cell centroid (**Fig. S1A**). Custom code for these operations and the resulting windrose plot was written in Python v3.9.1 using scikit-image v0.19.3, pandas v1.4.4, SciPy v1.9.1, and numpy v1.23.2. ^92–94^ Code is available on Github: https://github.com/fritzlaylinlab/contractile-vacuole-pressure.

### Environmental perturbations and under agarose assays

To assess the effects of temperature and sorbitol concentration on *N. gruberi* cells (**Fig. 2A**), 1.5% agarose pads were prepared by microwaving low-melt agarose (Affymetrix, 32830 25 GM) in water or 50 mM sorbitol (Sigma, S1876-500G), and pipetting 1.5 mL of solution into wells of 6-well tissue culture treated plates (CellTreat, 229105). Agarose pads solidified at room temperature, and were then wrapped in parafilm and stored at 4 °C for up to 1 month. Prior to the start of an experiment, agarose pads were warmed to room temperature. To control the temperature, an air conditioning unit in the microscope room was turned on >1 h prior to the experiment to achieve a stable temperature of 20±1 °C for the colder condition, and a stage heater (okolab UNO stage top incubator) was set to 35 °C for the warmer condition. 3X10^5^ - 5X10^5^ *Naegleria gruberi* NEG-M cells (grown at room temperature overnight to prevent differentiation to flagellates) were seeded into each well of a fresh 6-well tissue culture plate and allowed to settle for ~5 min. For each well, media was aspirated, cells were rinsed with 2 mL of water, and then 1 mL of water or 50 mM sorbitol was added. Using a small scoopula, agarose pads with matching concentrations of sorbitol were loosened at the edges, and a wedge was removed, resulting in a pac-man shape. The agarose pad was transferred onto the cells, and the solution was pipetted out of the well from the mouth of the pac-man. Sample preparation was staggered such that each well was treated approximately 1 h before imaging. Cells were transitioned to the proper temperature at least 20 min prior to imaging, and imaged for 16 min with 2 s between images using phase contrast. Experiments shown in **Fig. 2B** were performed similarly at room temperature (~24 °C), with the following exceptions: after seeding, cells were rinsed in 1 mL of 0-100 mM sorbitol, then 0.5 mL of 0-100 mM sorbitol were added prior to the addition of the agarose pad; cells were imaged after ~30 minutes of treatment; cells were imaged for 10 min total.

For **Fig. 2A-B**, conditions were tested in a different order for each of 5 trials (although the 20 °C condition was always completed before the 35 °C condition), and file names were blinded prior to analysis. For each replicate, 10 cells (**Fig. 2A**) or 5 cells (**Fig. 2B**) from each condition were randomly selected from the initial field of view for analysis (cells were only reselected if they left the field of view or divided during the movie). The largest recurring vacuole in each cell (the “main contractile vacuole”) was analyzed by measuring its area at its largest point prior to the pump using the freehand selection tool and measure command in Fiji. The pumping rate (frequency) was calculated by dividing the number of pumps −1 by the time between the first collapse and the last fullest point. Data were analyzed and graphs were created using GraphPad Prism.

### Comparisons of N. gruberi and D. discoideum under agarose

For the side-by-side comparisons of *N. gruberi* and *D. discoideum* (**Fig. 2C-D, Fig. S2**), 1.0% agarose pads in 12.5-100 mM sorbitol were prepared and stored as described above. Approximately 5X10^5^ *Naegleria gruberi* NEG-M cells (grown at room temperature overnight to prevent differentiation to flagellates) or 1X10^6^ *Dictyostelium discoideum* Ax2 cells were seeded into each well of a 6 well plate and allowed to settle. Media was aspirated from cells, and 1 mL of fresh media was added to each well prior to starting the experiment. For each well, media was aspirated, cells were rinsed with 2 mL followed by 1 mL of water or 12.5-100 mM sorbitol, and the agarose pad was transferred onto the cells as described above. 10 minutes were intentionally left between wells, to maintain a consistent timing of approximately 1 hour between exposure to water/sorbitol and imaging. Cells were imaged for 8 min with 2 s between images using phase contrast. Conditions were tested in a different order for each trial, and file names were blinded prior to analysis. For each replicate, 5 cells from each condition were randomly selected from the initial field of view for analysis (cells were only reselected if they left the field of view or divided during the movie). Every time a vacuole pumped (i.e. disappeared), the vacuole was measured at its largest point prior to the pump using the freehand selection and measure command in Fiji. Because every pumping event was measured (not only the main contractile vacuole as in **Fig. 2A-B**), the pumping rate was calculated by dividing the total number of pumps by the length of the movie (8 min). Data were analyzed and graphs were created using GraphPad Prism.

### Chemical inhibitors

Chemical inhibitor experiments (**Fig. 3B**) were performed under agarose at room temperature as described above (see: Environmental perturbations under agarose), with the following modifications. When making 1.5% agarose pads, small molecule inhibitors and controls were incorporated for final concentrations of: 5 μM Latrunculin B (Abcam, ab144291); 50 μM CK-666 (Calbiochem/Sigma, 182515); 50 μM CK-689 (Calbiochem/Sigma, 182517); or 0.1% DMSO (Sigma, D2650-5×5ML). After media was aspirated from cells, 1 mL of water was used to rinse cells, and 1 mL of inhibitor or control solutions (diluted in water) was added. Cells were imaged by phase contrast, beginning ~20 minutes after the addition of the agarose pad, and for 15 minutes with 2 seconds between frames. 20 minutes were left between wells to stagger the imaging times. Images of *D. discoideum* were taken at a rate of 250 ms/frame. Cells were imaged under 1.5% agarose pads with inhibitors as described above (or using Nocodazole (Sigma, M1404) at 10 μg/ml). Experiments with Bafilomycin (**Fig. 1D, Fig. S1D**) were performed in a similar way with the following exceptions: 100 nM Bafilomycin A1 (Sigma, 19-148), 0.1% DMSO (Sigma, D2650-5×5ML), or 0.1% water were used in the 1.5% agarose pads and to make the inhibitor or control solutions; agarose pads and solutions were diluted into 10% M7 media instead of water; and cells were seeded into 6-well glass-bottom plates (Cellvis, P06-1.5H-N).

Actin staining of DMSO and Latrunculin-treated *N. gruberi* cells (**Fig. S3A**) was performed live on the microscope using the OneStep fixing and staining protocol.^95^ Briefly, ~2X10^4^ cells (in 150 μl of M7 media) were seeded into a 96-well glass bottom plate (dot scientific, MGB096-1-2-LG-L), and imaged using DIC. During imaging, cells were treated with 150 μl of 0.2% DMSO or 10 μM LatB (for final concentrations of 0.1% DMSO and 5 μM LatB). We verified that DMSO- and Latrunculin-treated cells continued to pump their contractile vacuoles. Then, 150 μl of solution was removed from the well and 150 μl of OneStep fixing and staining solution (100 mM sucrose, 50 mM sodium phosphate buffer, 3.6% PFA, 0.025% NP-40, DAPI, and Alexa Fluor 488 Phalloidin-488: see ^95^ for additional details) were added. Imaging continued, adding in fluorescence to detect actin polymer and DNA. Once the staining had plateaued, a Z-stack was taken to encompass the thickness of the cells (step size: 0.6 μm, 40 slices). The images shown in **Fig. S3A** are maximum intensity projections.

### Microscopy

The experiments shown in **Fig. 1B**, **Fig. 2A**, and **Fig. S3A** were performed using a Nikon eclipse Ti2, equipped with a Crest X-Light spinning disk (50 μm pinhole), a Photometrics Prime 95B camera, a Lumencor Celesta light source for fluorescence, controlled through NIS-Elements AR software. Images in **Fig. 1B** and **Fig. S3A** were taken in DIC (acquired using transmitted light) and Fluorescence (excitation wavelengths: 546 (FM4-64), 477 (Alexa fluor 488 Phalloidin), and 405 (DAPI); emission wavelengths: 620 (FM4-64), 535 (Alexa fluor 488 Phalloidin), and 460 (DAPI)) with a Plan Apo λ 100x 1.45 NA oil objective (**Fig. 1B**) or a Plan Apo λ 40x 0.95 NA air objective (**Fig. S3A**). Experiments shown in **Fig. 2A** were performed without the spinning disk, using a ph2 40x S Plan Fluor 0.6 NA objective (with a correction collar set to 1.2).

The images in **Fig. 1C-D**, **Fig. 2B-D**, **Fig. 3B**, **Fig. 4B**, and **Fig. S1A** were collected using a Nikon eclipse Ti2, equipped with a pco.panda sCMOS camera, external phase system, and controlled through NIS-Elements AR software. A Plan Apo λ 100x 1.45 NA oil objective was used for **Fig. 1C** and **Fig. 4B**, a Plan Fluor 40x 1.3 NA oil objective was used for **Fig. 1D, Fig. S1A,** and **Fig. S1D**, and a ph2 40x S Plan Fluor 0.6 NA air objective (with a correction collar set to 1.2) was used for **Fig. 2B-D, Fig, 3B**.

Microscopy for **Fig. 4C** was completed using Nikon eclipse Ti2 equipped with a Ds-QI2 cMOS camera, external phase, and Plan Apo 40x 1.2 NA oil objective, controlled through NIS-Elements AR software. Experiments in **Fig. 3D** and **Fig. S3C** were completed using a Zeiss 980 laser scanning confocal with AiryScan2 equipped with a Plan Apo 63x 1.4 NA oil objective, and controlled by Zen Blue software.

### Line scans

The line scans shown in **Fig. 3D** and **Fig. S5B** were generated by drawing a line through a contractile vacuole and using the plot profile tool in Fiji. The same line was used for multiple frames, to determine the intensity along the line over time. The data were exported into GraphPad Prism to plot final graphs. **Fig. 3D** used a line thickness of 5 pixels, those in **Fig. S5B** used a line thickness of 10 pixels.

### Quick-Freeze Deep-Etch Electron Microscopy

Amoebae were scraped from bacteria-seeded agar plates and quick-frozen using the Heuser-designed slammer/cryopress device.^96^, ^97^ The freezing, which occurs at liquid helium (−269°C, 4°K) temperatures, is sufficiently rapid that water is immobilized in its vitreous state and does not associate to form distorting ice crystals. The frozen samples were stored at −196°C (liquid N_2_) and transferred to an evacuated Balzers BAF 400 D Freeze Etching System (Balzers AG, Lichtenstein), cooled with liquid N_2_, for fracturing. Fracturing was followed by etching, during which the temperature is raised from −196°C to −100°C for two minutes, allowing a surface layer of frozen water to sublimate into the vacuum, and leaving behind cellular elements and macromolecules suspended in their physiological state. The temperature was then reduced, and a film of vaporized platinum (Pt) was deposited onto the fractured surface from an angle of 25°. The sample is rapidly rotated during the deposition process and hence exposed to the Pt stream from all directions. A thin film of carbon was then deposited to hold the Pt grains in place. Pt does not condense on the surface as a uniform and amorphous coat. Instead it forms small crystals, 10-20 Å in diameter, which migrate and clump together in various ways depending on the texture and composition of the surface. Hence the resultant replica is a high-resolution platinum cast which mimics the contours of the exposed sample. The replica was cleaned with chromosulfuric acid to remove biological material, generating a very thin film of Pt/C – the replica -- that was placed on a formvar-coated copper grid and viewed using the electron beam of a transmission electron microscope (JEOL model JEM 1400 (JEOL, Peabody, MA 01960)), equipped with an AMT (Advanced Microscopy Techniques, Woburn, MA 01801, USA) V601 digital camera.

### Biophysical modeling of contractile vacuole emptying

The cytosolic pressure model assumes that all of the contents of the vacuole exit through a single pore in the membrane. The pore is assumed to have constant radius and thickness throughout time. The fluid in the vacuole is assumed to be an incompressible Newtonian fluid (water) engaging in laminar flow through the pore. The flow through this pore is therefore determined by the Hagen–Poiseuille law for laminar flow through a cylinder: 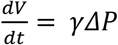. The pressure difference across the pore (i.e., between the inside of the vacuole and the extracellular space) forces fluid through the pore. The small size of the pore gives a resistance to fluid flow, defined by *γ*. For a cylindrical pore, 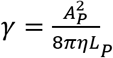, where A_P_ and L_P_ are the area and length of the pore, respectively, and η is the viscosity of the medium (**Fig. 4A**). See the **Text S2** for a full derivation for the equation shown in **Fig. 4A**.

Importantly, this model assumes the membrane is impermeable to water at short timescales, such that water can only leave through the pore. Water cannot leave or enter through the membrane itself. We can make this approximation because the rate of fluid flow through the vacuole membrane (e.g., the vacuole fill rate) is more than ten times slower than that of the fluid expulsion rate through the pore.

### Statistical analysis

Experiments shown in **Fig. 1, 2, 3B** and **3C** were each performed at least 5 times. All data were analyzed using Graphpad Prism, and plotted as SuperPlots^98^ when possible. Because all comparisons included more than two conditions, statistical significance between experiment-level averages was determined using ordinary one-way ANOVAs with Tukey’s multiple comparisons tests to calculate p values. 0.05 was used as a cutoff for significance.

## ACKNOWLEDGEMENTS

We thank Dr. Ursula Goodenough (Washington University) for providing Quick-Freeze Deep-Etch Electron Microscopy images of *Naegleria* cells, Dr. Chandler Fulton (Brandeis University) for providing NEG cells for Quick-Freeze Deep-Etch Electron Microscopy and the NEG-M cell line. We thank Dr. Nicholas Martin (University of California San Francisco) and Manuela Richter (University of California San Francisco) for important discussions and experiments during the Physiology Course at the Marine Biological Laboratory. We also thank Erik Kalinka for translating key sections of the reference (Schardinger, 1899) into English, and Dr. Sam Lord, Dr. Ken Campellone, Dr. Madelaine Bartlett, and Dr. Meg Titus for providing feedback on this work. This work was supported by: the National Institute Of Allergy And Infectious Diseases of the National Institutes of Health under Award Number F32AI150057 to K.B.V.; the National Institute of General Medical Science of the National Institutes of Health under award number K99GM147656 to K.B.V., award number F32GM148023 to A.S.K., award number R35GM143039 to L.K.F.-L., and award number 1R15GM143733-01 to M.E.; a Smith Family Foundation Award for Excellence in Biomedical Science to L.K.F.-L.; and The Pew Charitable Trusts (award to L.K.F.-L.). R.M.G and C.L. were supported by The National Science Foundation Simons Center for Mathematical and Statistical Analysis of Biology at Harvard (Award #1764269). L.K.F.-L. Is a fellow of the Canadian Institute for Advanced Research Fungal Kingdom Program.

## SUPPLEMENTAL ITEMS

**Figure S1. Characterizing *Naegleria* contractile vacuoles and their mechanisms.** (**A**) (*Left*) A representative *N. gruberi* amoeba under agarose illustrates the definition of the orientation angle, *θ*, between the centroid velocity and the contractile vacuole. (*Right*) A windrose plot shows the orientation of the contractile vacuole at the time of pumping relative to the velocity of the cell’s centroid at that moment. The histogram depicts 80 pumping events from 15 cells across 5 biological replicates. Open colored circles show the average orientation from populations of cells measured on 5 separate days (2-4 cells and 13-19 pumps per replicate); the gray square shows the average of these 5 biological replicates. (**B**) Alignments produced by T-COFFEE of the hits for V_1_ subunit G, show the sequence similarity of these genes. Residues are colored to indicate amino acids with similar properties, with darker shading indicating greater conservation. (**C**) Alignments of V_0_ subunits *c* and *a*. Key residues are highlighted, including those that form the Bafilomycin A binding site (yellow) or are functionally disrupted by Bafilomycin A binding (cyan). Residues are based on cryo-EM structures of the *B. taurus* v-ATPase.^99^ (**D**) Cells were incubated with or without the vacuolar-type H+ ATPase inhibitor Bafilomycin A1 or DMSO (carrier control). Each contractile vacuole bladder was measured at its maximum size, and the cumulative contractile vacuole area was calculated over 8 min. Left panels show the cumulative area pumped out of the cell for 25 cells: 5 cells each from 5 experiments. The right panel shows the area pumped per minute, with each small gray symbol representing a single cell (5 cells per experiment), and larger symbols representing experiment-level averages for 5 replicates, with symbols coordinated by experiment.

**Figure S2. *Naegleria* and *Dictyostelium* contractile vacuole bladders are larger under more hypo-osmotic stress.** Violin plots show the maximum contractile vacuole area for each pumping event in *N. gruberi* (**A**) or *D. discoideum* (**B**) for each of five experimental replicates, which were pooled in **Fig. 2C-D**. White lines indicate the median and quartiles.

**Figure S3. *Naegleria* and *Dictyostelium* contractile vacuole emptying can occur in the absence of cytoskeletal polymers.** (**A**) *N. gruberi* cells were treated with DMSO or Latrunculin B, then fixed and stained for actin polymer (Alexa-488 conjugated phalloidin) and DNA (DAPI). Maximum intensity projections of representative cells are shown, adjusted to the same brightness and contrast settings. (**B**) The same DMSO and Latrunculin data shown in **Fig. 3C** is shown within the context of a wider drug panel (including the Arp2/3 complex inhibitor CK-666 and the microtubule inhibitor Nocodazole). Graphs show the area pumped per minute by *Dictyostelium* contractile vacuoles, the number of pumps per minute, and the maximum contractile vacuole area per cell. (**C**) A *D. discoideum* cell expressing a fluorescently-labeled actin-binding protein is shown through a pumping cycle. Line scans bisecting a contractile vacuole show no enrichment of actin around the pumping bladder.

**Figure S4. Estimating contractile vacuole pore size.** (**A**) Model parameters for the cytoplasmic pressure model that allow the contractile vacuole to dispense its entire volume in 1 s. The required pore diameter is plotted as a function of the choice of pore width, given a cytoplasmic pressure of 10 Pa (solid line), 100 Pa (dashed line), or 1000 Pa (dotted line). Any reasonable choice of cytoplasmic pressure in between 10 and 1000 Pa (gray shading) allows for a wide range of possible pore diameters and pore depths. (**B**) A table shows the allowable parameters^66^ representing the results shown in (A).

**Video S1. Contractile vacuole networks in *Naegleria* comprise small vacuoles and large bladders, with multiple generations co-existing.** (**A**) An *N. gruberi* cell stained with the membrane dye FM4-64 cell shows a contractile vacuole pumping cycle. Frames from this movie were selected for **Fig. 1B.** (**B**) A representative cell shows multiple contractile vacuole pumping cycles, with full bladders pseudocolored through their expulsion. Note the surrounding network of uncolored vacuoles grows as each pseudocolored bladder pumps. This movie was also used in **Fig. 1C**.

**Video S2. *Naegleria* contractile vacuole networks are dynamic.** (**A**) A representative contractile vacuole fusion event is shown, with initial contractile vacuoles fusing to form a single bladder at 3 min 34 s. The two networks that merge are indicated with pink asterisks, displayed for 2 frames prior to the start of the pump. The single merged bladder is indicated with a magenta asterisk. (**B**) A representative fission event is shown, where a single bladder (magenta asterisk) becomes smaller contractile vacuoles (green asterisks). (**C**) A cell with multiple large and small contractile vacuole networks is shown. The two largest systems are indicated with yellow and cyan asterisks. Cells in A and B were untreated control cells from the experiments shown in **Fig. S1D**. Cells in C are from a control experiment shown in **Fig. 3B** (DMSO-treated control).

**Video S3. Latrunculin does not prevent contractile vacuole pumping events in *Naegleria*.** A representative video of LatB-treated *N. gruberi* cells shows that contractile vacuole expulsion events are still common.

**Video S4. Actin does not localize to *Dictyostelium* contractile vacuoles.** A representative video of an RFP-LimE-expressing *D. discoideum* cell (the same example shown in **Fig. 3D**) shows no enrichment of actin at contractile vacuoles (highlighted with green asterisks).

**Text S1. Modeling contractile vacuole filling Text S2. Modeling contractile vacuole emptying**

## REFERENCES

1. Schardinger, F. (1899). Entwicklungskreis eine Amoeba lobosa (gymnamoeba): Amoeba gruberi. Sitzungsberichte der Akademie der Wissenschaften mathematisch-naturwissenschaftliche Klasse 108, 713–734.

2. Coppellotti, O., Piccinni, E., Colombetti, G., and Leno, F. (1979). Responses of Euglena gracilis to Cytochalasins B and D. Italian Journal of Zoology 46, 71–75.

3. Figarella, K., Uzcategui, N.L., Zhou, Y., LeFurgey, A., Ouellette, M., Bhattacharjee, H., and Mukhopadhyay, R. (2007). Biochemical characterization of Leishmania major aquaglyceroporin LmAQP1: possible role in volume regulation and osmotaxis. Mol Microbiol 65, 1006–1017.

4. Jimenez, V., Miranda, K., and Augusto, I. (2022). The old and the new about the contractile vacuole of Trypanosoma cruzi. J Eukaryot Microbiol, e12939.

5. Troster, V., Setzer, T., Hirth, T., Pecina, A., Kortekamp, A., and Nick, P. (2017). Probing the contractile vacuole as Achilles’ heel of the biotrophic grapevine pathogen Plasmopara viticola. Protoplasma 254, 1887–1901.

6. Naitoh, Y., Tominaga, T., Ishida, M., Fok, A., Aihara, M., and Allen, R. (1997). How does the contractile vacuole of Paramecium multimicronucleatum expel fluid? Modelling the expulsion mechanism. J Exp Biol 200, 713–721.

7. Komsic-Buchmann, K., Wostehoff, L., and Becker, B. (2014). The contractile vacuole as a key regulator of cellular water flow in Chlamydomonas reinhardtii. Eukaryot Cell 13, 1421–1430.

8. Toret, C., Picco, A., Boiero-Sanders, M., Michelot, A., and Kaksonen, M. (2022). The cellular slime mold Fonticula alba forms a dynamic, multicellular collective while feeding on bacteria. Curr Biol.

9. Gabriel, D., Hacker, U., Köhler, J., Müller-Taubenberger, A., Schwartz, J.-M., Westphal, M., and Gerisch, G. (1999). The contractile vacuole network of Dictyostelium as a distinct organelle: its dynamics visualized by a GFP marker protein. J Cell Science.

10. Doberstein, S.K., Baines, I.C., Wiegand, G., Korn, E.D., and Pollard, T.D. (1993). Inhibition of contractile vacuole function in vivo by antibodies against myosin-I. Nature 365, 841–843.

11. Wigg, D., Bovee, E.C., and Jahn, T.L. (1967). The evacuation mechanism of the water expulsion vesicle (“contractile vacuole”) of Amoeba proteus. J Protozool 14, 104–108.

12. Spallanzani, L., and Bonnet, C. (1776). Opuscoli di fisica animale, e vegetabile dell’Abate Spallanzani, Regio Professore di Storia Naturale nell’Università di Pavia, socio delle Accademie di Londra de’Curiosi della Natura di Germania, di Berlino, Stockolm, Gottinga, Bologna, Siena, ec.: aggiuntevi alcune lettere relative, (In Modena: Presso la Società tipografica).

13. Martin-Navarro, C.M., Lorenzo-Morales, J., Lopez-Arencibia, A., Reyes-Batlle, M., Pinero, J.E., Valladares, B., and Maciver, S.K. (2014). Evaluation of Acanthamoeba myosin-IC as a potential therapeutic target. Antimicrobial Agents and Chemotherapy 58, 2150–2155.

14. Becker, B., and Hickisch, A. (2005). Inhibition of contractile vacuole function by brefeldin A. Plant Cell Physiol 46, 201–212.

15. Clarke, M., Kohler, J., Arana, Q., Liu, T., Heuser, J., and Gerisch, G. (2002). Dynamics of the vacuolar H(+)-ATPase in the contractile vacuole complex and the endosomal pathway of Dictyostelium cells. Journal of Cell Science 115, 2893–2905.

16. Heuser, J., Zhu, Q., and Clarke, M. (1993). Proton pumps populate the contractile vacuoles of Dictyostelium amoebae. J Cell Biol 121, 311–327.

17. Ishida, M., Hori, M., Ooba, Y., Kinoshita, M., Matsutani, T., Naito, M., Hagimoto, T., Miyazaki, K., Ueda, S., Miura, K., et al. (2021). A functional Aqp1 gene product localizes on the contractile vacuole complex in Paramecium multimicronucleatum. J Eukaryot Microbiol 68, e12843.

18. Montalvetti, A., Rohloff, P., and Docampo, R. (2004). A functional aquaporin co-localizes with the vacuolar proton pyrophosphatase to acidocalcisomes and the contractile vacuole complex of Trypanosoma cruzi. J Biol Chem 279, 38673–38682.

19. Nishihara, E., Yokota, E., Tazaki, A., Orii, H., Katsuhara, M., Kataoka, K., Igarashi, H., Moriyama, Y., Shimmen, T., and Sonobe, S. (2008). Presence of aquaporin and V-ATPase on the contractile vacuole of Amoeba proteus. Biology of the Cell 100, 179–188.

20. Ruiz, F.A., Marchesini, N., Seufferheld, M., Govindjee, and Docampo, R. (2001). The polyphosphate bodies of Chlamydomonas reinhardtii possess a proton-pumping pyrophosphatase and are similar to acidocalcisomes. J Biol Chem 276, 46196–46203.

21. Stock, C., Gronlien, H.K., Allen, R.D., and Naitoh, Y. (2002). Osmoregulation in Paramecium: in situ ion gradients permit water to cascade through the cytosol to the contractile vacuole. Journal of Cell Science 115, 2339–2348.

22. Ulrich, P.N., Jimenez, V., Park, M., Martins, V.P., Atwood, J., 3rd, Moles, K., Collins, D., Rohloff, P., Tarleton, R., Moreno, S.N., et al. (2011). Identification of contractile vacuole proteins in Trypanosoma cruzi. PLoS One 6, e18013.

23. Young, R.A. (1924). On the excretory apparatus in Paramecium. Science 60, 244.

24. Tani, T., Tominaga, T., Allen, R.D., and Naitoh, Y. (2002). Development of periodic tension in the contractile vacuole complex membrane of paramecium governs its membrane dynamics. Cell Biology International 26, 853–860.

25. Gerisch, G., Heuser, J., and Clarke, M. (2002). Tubular-vesicular transformation in the contractile vacuole system of Dictyostelium. Cell Biology International 26, 845–852.

26. Nishihara, E., Shimmen, T., and Sonobe, S. (2007). New aspects of membrane dynamics of Amoeba proteus contractile vacuole revealed by vital staining with FM 4-64. Protoplasma 231, 25–30.

27. McKanna, J.A. (1973). Fine structure of the contractile vacuole pore in Paramecium. J Protozool 20, 631–638.

28. Heuser, J. (2006). Evidence for recycling of contractile vacuole membrane during osmoregulation in Dictyostelium amoebae--a tribute to Gunther Gerisch. Eur J Cell Biol 85, 859–871.

29. Gerald, N.J., Siano, M., and De Lozanne, A. (2002). The Dictyostelium LvsA protein is localized on the contractile vacuole and is required for osmoregulation. Traffic 3, 50–60.

30. Docampo, R., Jimenez, V., Lander, N., Li, Z.H., and Niyogi, S. (2013). New insights into roles of acidocalcisomes and contractile vacuole complex in osmoregulation in protists. Int Rev Cell Mol Biol 305, 69–113.

31. Heath, R.J., and Insall, R.H. (2008). Dictyostelium MEGAPs: F-BAR domain proteins that regulate motility and membrane tubulation in contractile vacuoles. Journal of Cell Science 121, 1054–1064.

32. Baines, I.C., and Korn, E.D. (1990). Localization of myosin IC and myosin II in Acanthamoeba castellanii by indirect immunofluorescence and immunogold electron microscopy. Journal of Cell Biology 111, 1895–1904.

33. Baines, I.C., Corigliano-Murphy, A., and Korn, E.D. (1995). Quantification and localization of phosphorylated myosin I isoforms in Acanthamoeba castellanii. Journal of Cell Biology 130, 591–603.

34. Yonemura, S., and Pollard, T. (1992). The localization of myosin I and myosin II in Acanthamoeba by fluorescence microscopy. J Cell Science 102, 629–642.

35. Kong, H.H., and Pollard, T.D. (2002). Intracellular localization and dynamics of myosin-II and myosin-IC in live Acanthamoeba by transient transfection of EGFP fusion proteins. Journal of Cell Science 115, 4993–5002.

36. Bement, W.M., and Mooseker, M.S. (1993). Keeping out the rain. Nature 365, 785–786.

37. Betts, H.C., Puttick, M.N., Clark, J.W., Williams, T.A., Donoghue, P.C.J., and Pisani, D. (2018). Integrated genomic and fossil evidence illuminates life’s early evolution and eukaryote origin. Nat Ecol Evol 2, 1556–1562.

38. Knoll, A.H. (2014). Paleobiological perspectives on early eukaryotic evolution. Cold Spring Harb Perspect Biol 6.

39. Parfrey, L.W., Lahr, D.J., Knoll, A.H., and Katz, L.A. (2011). Estimating the timing of early eukaryotic diversification with multigene molecular clocks. Proc Natl Acad Sci U S A 108, 13624–13629.

40. Fulton, C., and Dingle, A.D. (1971). Basal bodies, but not centrioles, in Naegleria. J Cell Biol 51, 826–836.

41. Lee, J.H., and Walsh, C.J. (1988). Transcriptional regulation of coordinate changes in flagellar mRNAs during differentiation of Naegleria gruberi amebae into flagellates. Mol Cell Biol 8, 2280–2287.

42. Chung, S., Cho, J., Cheon, H., Paik, S., and Lee, J. (2002). Cloning and characterization of a divergent alpha-tubulin that is expressed specifically in dividing amebae of Naegleria gruberi. Gene 298, 77–86.

43. Walsh, C.J. (2007). The role of actin, actomyosin and microtubules in defining cell shape during the differentiation of Naegleria amebae into flagellates. Eur J Cell Biol 86, 85–98.

44. Walsh, C.J. (2012). The structure of the mitotic spindle and nucleolus during mitosis in the amebo-flagellate Naegleria. PLoS One 7, e34763.

45. Velle, K.B., Kennard, A.S., Trupinic, M., Ivec, A., Swafford, A.J.M., Nolton, E., Rice, L.M., Tolic, I.M., Fritz-Laylin, L.K., and Wadsworth, P. (2022). Naegleria’s mitotic spindles are built from unique tubulins and highlight core spindle features. Curr Biol.

46. Jung, G., Titus, M.A., and Hammer, J.A., 3rd (2009). The Dictyostelium type V myosin MyoJ is responsible for the cortical association and motility of contractile vacuole membranes. Journal of Cell Biology 186, 555–570.

47. Wilson, C.W. (1916). On the life history of a soil amoeba. University of California Press, 241–292.

48. Siddiqui, R., Ali, I.K.M., Cope, J.R., and Khan, N.A. (2016). Biology and pathogenesis of Naegleria fowleri. Acta Trop 164, 375–394.

49. Goldberg, N.B., Sawinski, V.J., and Goldberg, A.F. (1965). Human cerebrospinal fluid osmolality at 37-degree C. Anesthesiology 26, 829.

50. Albers, T., Maniak, M., Beitz, E., and von Bülow, J. (2016). The C isoform of Dictyostelium tetraspanins localizes to the contractile vacuole and contributes to resistance against osmotic stress. PLoS One 11.

51. Becker, M., Matzner, M., and Gerisch, G. (1999). Drainin required for membrane fusion of the contractile vacuole in Dictyostelium is the prototype of a protein family also represented in man. EMBO J 18, 3305–3316.

52. Clarke, M., Maddera, L., Engel, U., and Gerisch, G. (2010). Retrieval of the vacuolar H-ATPase from phagosomes revealed by live cell imaging. PLoS One 5, e8585.

53. Du, F., Edwards, K., Shen, Z., Sun, B., De Lozanne, A., Briggs, S., and Firtel, R.A. (2008). Regulation of contractile vacuole formation and activity in Dictyostelium. EMBO J 27, 2064–2076.

54. Essid, M., Gopaldass, N., Yoshida, K., Merrifield, C., and Soldati, T. (2012). Rab8a regulates the exocyst-mediated kiss-and-run discharge of the Dictyostelium contractile vacuole. Mol Biol Cell 23, 1267–1282.

55. Malchow, D., Lusche, D.F., Schlatterer, C., De Lozanne, A., and Muller-Taubenberger, A. (2006). The contractile vacuole in Ca2+-regulation in Dictyostelium: its essential function for cAMP-induced Ca2+-influx. BMC Developmental Biology 6, 31.

56. Manna, P.T., Barlow, L.D., Ramirez-Macias, I., Herman, E.K., and Dacks, J.B. (2023). Endosomal vesicle fusion machinery is involved with the contractile vacuole in Dictyostelium discoideum. Journal of Cell Science 136.

57. Maringer, K., Yarbrough, A., Sims-Lucas, S., Saheb, E., Jawed, S., and Bush, J. (2016). Dictyostelium discoideum RabS and Rab2 colocalize with the Golgi and contractile vacuole system and regulate osmoregulation. J Biosci 41, 205–217.

58. Nolta, K.V., and Steck, T.L. (1994). Isolation and initial characterization of the bipartite contractile vacuole complex from Dictyostelium discoideum. J Biol Chem 269, 2225–2233.

59. Zhu, Q., and Clarke, M. (1992). Association of calmodulin and an unconventional myosin with the contractile vacuole complex of Dictyostelium discoideum. Journal of Cell Biology 118, 347–358.

60. Page, F.C. (1967). Taxonomic Criteria for Limax Amoebae, with Descriptions of 3 New Species of Hartmannella and 3 of Vahlkampfia. J Protozool 14, 499–521.

61. Fok, A.K., Aihara, M.S., Ishida, M., Nolta, K.V., Steck, T.L., and Allen, R.D. (1995). The pegs on the decorated tubules of the contractile vacuole complex of Paramecium are proton pumps. Journal of Cell Science 108 (Pt 10), 3163–3170.

62. Bowman, E.J., Siebers, A., and Altendork, K. (1988). Bafilomycins: A class of inhibitors of membrane ATPases from microorganisms, animal cells, and plant cells. Proc Natl Acad Sci U S A 85, 7972–7976.

63. Crider, B.P., Xie, X.S., and Stone, D.K. (1994). Bafilomycin inhibits proton flow through the H+ channel of vacuolar proton pumps. J Biol Chem 269, 17379–17381.

64. Dai, J., Ting-Beall, H.P., Hochmuth, R.M., Sheetz, M.P., and Titus, M.A. (1999). Myosin I contributes to the generation of resting cortical tension. Biophysical Journal 77, 1168–1176.

65. Manoussaki, D., Shin, W.D., Waterman, C.M., and Chadwick, R.S. (2015). Cytosolic pressure provides a propulsive force comparable to actin polymerization during lamellipod protrusion. Sci Rep 5, 12314.

66. Petrie, R.J., and Koo, H. (2014). Direct measurement of intracellular pressure. Curr Protoc Cell Biol 63, 12 19 11–19.

67. Cole, W.H. (1925). Pulsation of the contractile vacuole of Paramecium as affected by temperature.

68. Kitching, J.A. (1948). The physiology of contractile vacuoles; temperature and osmotic stress. J Exp Biol 25, 421–436.

69. Nematbakhsh, S., and Bergquist, B.L. (1993). Periodicity and the influence of temperature and cellular size in contractile vacuole contraction intervals. T Am Microsc Soc 112, 292–305.

70. Stock, C., Allen, R.D., and Naitoh, Y. (2001). How external osmolarity affects the activity of the contractile vacuole complex, the cytosolic osmolarity and the water permeability of the plasma membrane in Paramecium multimicronucleatum. J Exp Biol 204, 291–304.

71. Spector, I., Shochet, N.R., Kashman, Y., and Groweiss, A. (1983). Latrunculins: novel marine toxins that disrupt microfilament organization in cultured cells. Science 219, 493–495.

72. Velle, K.B., and Fritz-Laylin, L.K. (2020). Conserved actin machinery drives microtubule-independent motility and phagocytosis in Naegleria. Journal of Cell Biology 219.

73. Nolen, B.J., Tomasevic, N., Russell, A., Pierce, D.W., Jia, Z., McCormick, C.D., Hartman, J., Sakowicz, R., and Pollard, T.D. (2009). Characterization of two classes of small molecule inhibitors of Arp2/3 complex. Nature 460, 1031–1034.

74. Hetrick, B., Han, M.S., Helgeson, L.A., and Nolen, B.J. (2013). Small molecules CK-666 and CK-869 inhibit actin-related protein 2/3 complex by blocking an activating conformational change. Chem Biol 20, 701–712.

75. Gerisch, G., Bretschneider, T., Muller-Taubenberger, A., Simmeth, E., Ecke, M., Diez, S., and Anderson, K. (2004). Mobile actin clusters and traveling waves in cells recovering from actin depolymerization. Biophysical Journal 87, 3493–3503.

76. Mitchison, T.J. (2019). Colloid osmotic parameterization and measurement of subcellular crowding. Mol Biol Cell 30, 173–180.

77. Luykx, P., Hoppenrath, M., and Robinson, D.G. (1997). Structure and behavior of contractile vacuoles in Chlamydomonas reinhardtii. Protoplasma 198, 73–84.

78. Fulton, C. (1970). Amebo-flagellates as research partners: The laboratory biology of Naegleria and Tetramitus. In Methods in Cell Physiology, Volume 4. pp. 341–476.

79. Tani, T., Allen, R.D., and Naitoh, Y. (2000). Periodic tension development in the membrane of the in vitro contractile vacuole of Paramecium multimicronucleatum: modification by bisection, fusion and suction. J Exp Biol 203, 239–251.

80. Tani, T., Allen, R.D., and Naitoh, Y. (2001). Cellular membranes that undergo cyclic changes in tension: Direct measurement of force generation by an in vitro contractile vacuole of Paramecium multimicronucleatum. J Cell Sci 114, 785–795.

81. Bean, B.P. (2007). The action potential in mammalian central neurons. Nat Rev Neurosci 8, 451–465.

82. Lakatta, E.G., Maltsev, V.A., and Vinogradova, T.M. (2010). A coupled SYSTEM of intracellular Ca2+ clocks and surface membrane voltage clocks controls the timekeeping mechanism of the heart’s pacemaker. Circulation Research 106, 659–673.

83. Ren, D. (2011). Sodium leak channels in neuronal excitability and rhythmic behaviors. Neuron 72, 899–911.

84. Hellsten, M., and Roos, U.P. (1998). The actomyosin cytoskeleton of amoebae of the cellular slime molds Acrasis rosea and Protostelium mycophaga: structure, biochemical properties, and function. Fungal Genet Biol 24, 123–145.

85. Allen, R.D., and Fok, A.K. (1988). Membrane dynamics of the contractile vacuole complex of Paramecium. J Protozool 35, 63–71.

86. Cohen, J., Garreau de Loubresse, N., and Beisson, J. (1984). Actin microfilaments in Paramecium: localization and role in intracellular movements. Cell Motility 4, 443–468.

87. Pappas, G.D., and Brandt, P.W. (1958). The fine structure of the contractile vacuole in Ameba. J Biophys Biochem Cytol 4, 485–488.

88. Tominaga, T., Allen, R.D., and Naitoh, Y. (1998). Electrophysiology of the in situ contractile vacuole complex of Paramecium reveals its membrane dynamics and electrogenic site during osmoregulatory activity. J Exp Biol 201, 451–460.

89. Fey, P., Dodson, R.J., Basu, S., and Chisholm, R.L. (2013). One stop shop for everything Dictyostelium: dictyBase and the Dicty Stock Center in 2012. Methods Mol Biol 983, 59–92.

90. Gaudet, P., Fey, P., and Chisholm, R. (2008). Transformation of Dictyostelium with plasmid DNA by electroporation. CSH Protoc 2008, pdb prot5103.

91. Schindelin, J., Arganda-Carreras, I., Frise, E., Kaynig, V., Longair, M., Pietzsch, T., Preibisch, S., Rueden, C., Saalfeld, S., Schmid, B., et al. (2012). Fiji: an open-source platform for biological-image analysis. Nat Methods 9, 676–682.

92. McKinney, W. (2010). Data Structures for Statistical Computing in Python. Proceedings of the 9th Python in Science Conference 445.

93. van der Walt, S., Schonberger, J.L., Nunez-Iglesias, J., Boulogne, F., Warner, J.D., Yager, N., Gouillart, E., Yu, T., and scikit-image, c. (2014). scikit-image: image processing in Python. PeerJ 2, e453.

94. Harris, C.R., Millman, K.J., van der Walt, S.J., Gommers, R., Virtanen, P., Cournapeau, D., Wieser, E., Taylor, J., Berg, S., Smith, N.J., et al. (2020). Array programming with NumPy. Nature 585, 357–362.

95. Velle, K.B., Fermino do Rosario, C., Wadsworth, P., and Fritz-Laylin, L.K. (2021). A OneStep Solution to Fix and Stain Cells for Correlative Live and Fixed Microscopy. Curr Protoc 1, e308.

96. Heuser, J. (1981). Preparing biological samples for stereomicroscopy by the quick-freeze, deep-etch, rotary-replication technique. Methods Cell Biol 22, 97–122.

97. Heuser, J.E. (2011). The origins and evolution of freeze-etch electron microscopy. J Electron Microsc (Tokyo) 60 Suppl 1, S3–29.

98. Lord, S.J., Velle, K.B., Mullins, R.D., and Fritz-Laylin, L.K. (2020). SuperPlots: Communicating reproducibility and variability in cell biology. Journal of Cell Biology 219.

99. Wang, R., Wang, J., Hassan, A., Lee, C.H., Xie, X.S., and Li, X. (2021). Molecular basis of V-ATPase inhibition by bafilomycin A1. Nat Commun 12, 1782.

