## Supplement for "A conserved pressure-driven mechanism for regulating cytosolic osmolarity"

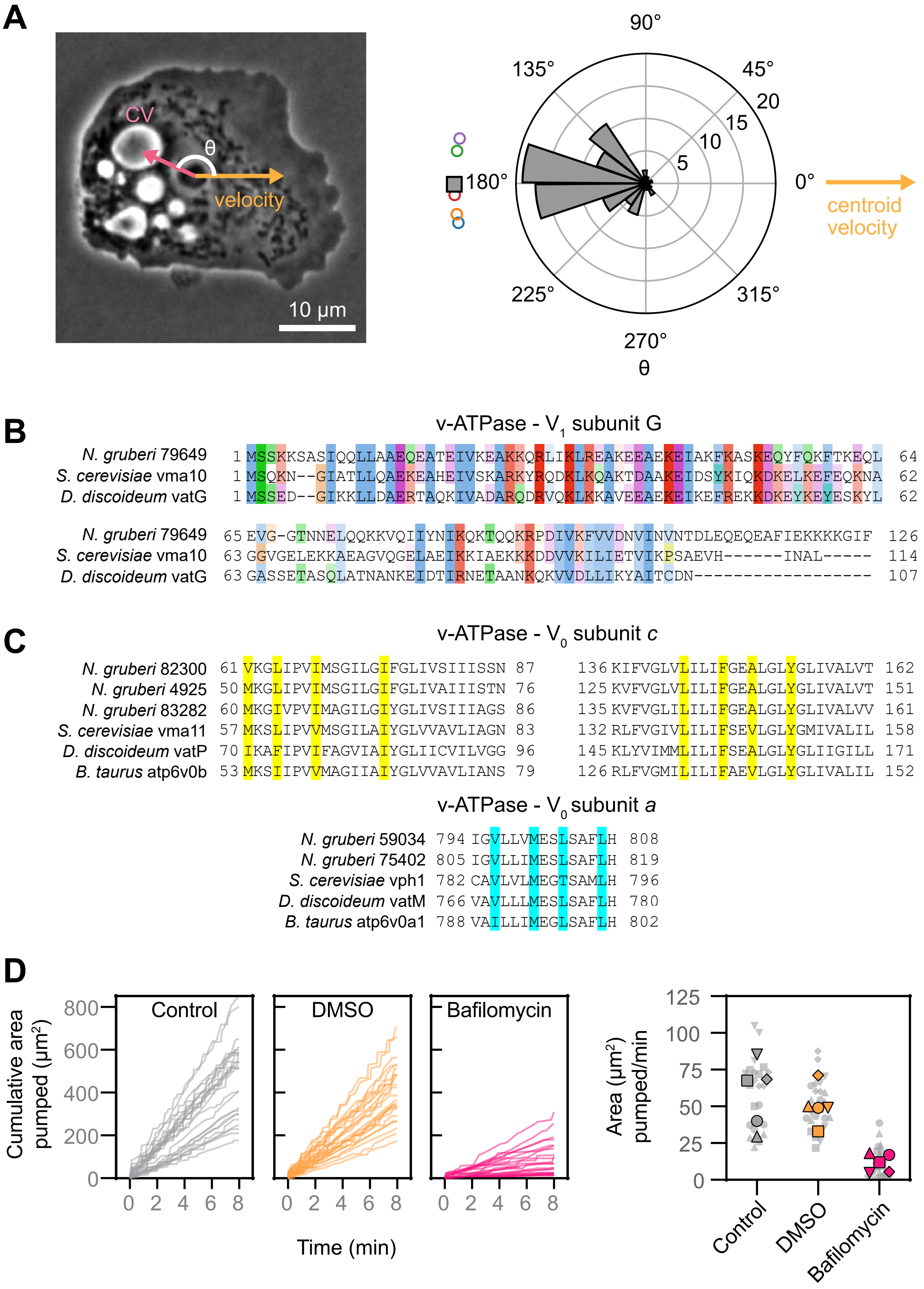

**Figure S1. Characterizing *Naegleria* contractile vacuoles and their mechanisms.** (**A**) *(Left)* A representative *N. gruberi* amoeba under agarose illustrates the definition of the orientation angle, *θ,* between the centroid velocity and the contractile vacuole. *(Right)* A windrose plot shows the orientation of the contractile vacuole at the time of pumping relative to the velocity of the cell's centroid at that moment. The histogram depicts 80 pumping events from 15 cells across 5 biological replicates. Open colored circles show the average orientation from populations of cells measured on 5 separate days (2-4 cells and 13-19 pumps per replicate); the gray square shows the average of these 5 biological replicates. (**B**) Alignments produced by T-COFFEE of the hits for $V_1$ subunit G, show the sequence similarity of these genes. Residues are colored to indicate amino acids with similar properties, with darker shading indicating greater conservation. (**C**) Alignments of $V_0$ subunits *c* and *a*. Key residues are highlighted, including those that form the Bafilomycin A binding site (yellow) or are functionally disrupted by Bafilomycin A binding (cyan). Residues are based on cryo-EM structures of the *B. taurus* v-ATPase.[99] (**D**) Cells were incubated with or without the vacuolar-type H+ ATPase inhibitor Bafilomycin A1 or DMSO (carrier control). Each contractile vacuole bladder was measured at its maximum size, and the cumulative contractile vacuole area was calculated over 8 min. Left panels show the cumulative area pumped out of the cell for 25 cells: 5 cells each from 5 experiments. The right panel shows the area pumped per minute, with each small gray symbol representing a single cell (5 cells per experiment), and larger symbols representing experiment-level averages for 5 replicates, with symbols coordinated by experiment.

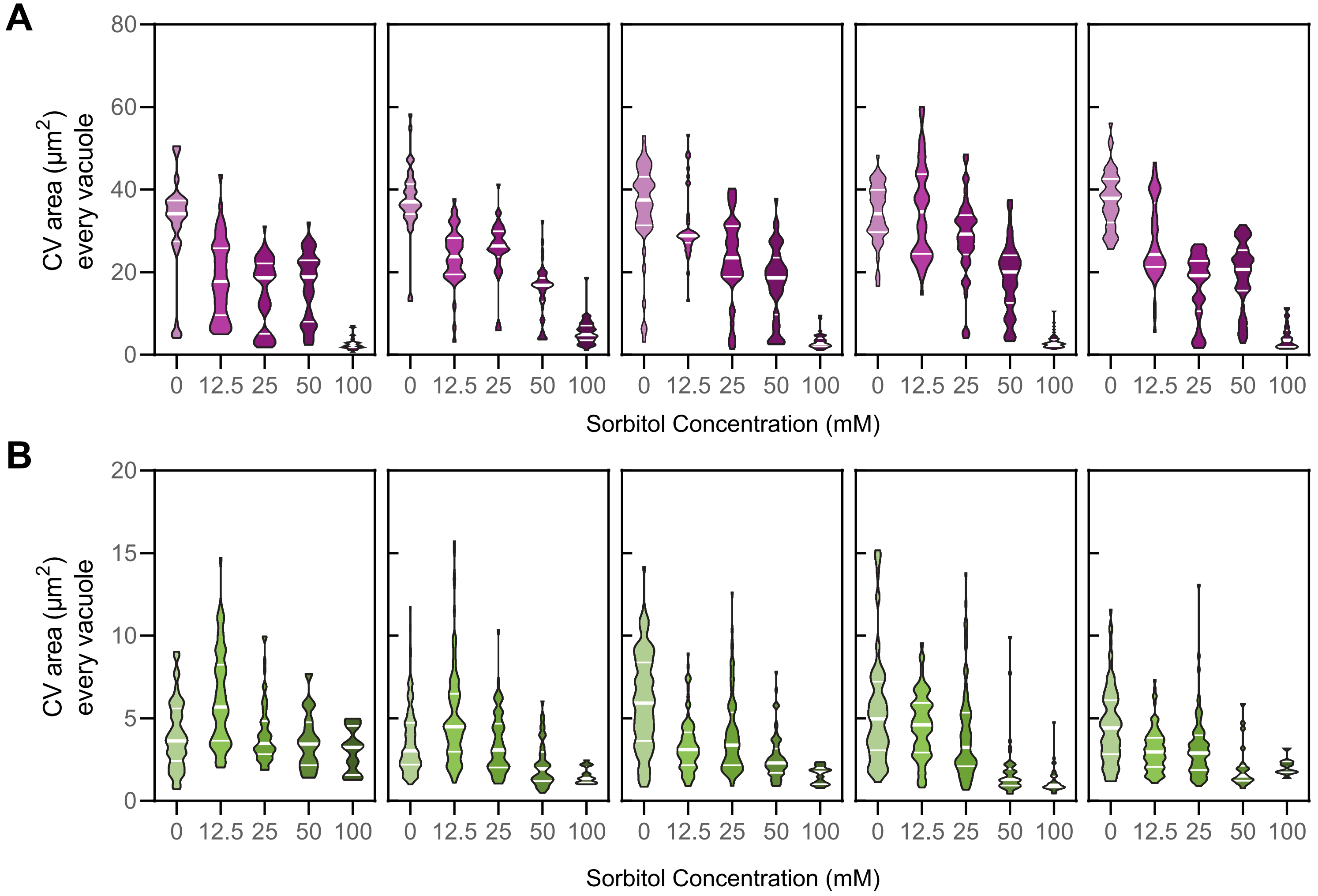

**Figure S2. *Naegleria* and *Dictyostelium* contractile vacuole bladders are larger under more hypo-osmotic stress.** Violin plots show the maximum contractile vacuole area for each pumping event in *N. gruberi* (**A**) or *D. discoideum* (**B**) for each of five experimental replicates, which were pooled in **Fig. 2C-D**. White lines indicate the median and quartiles.

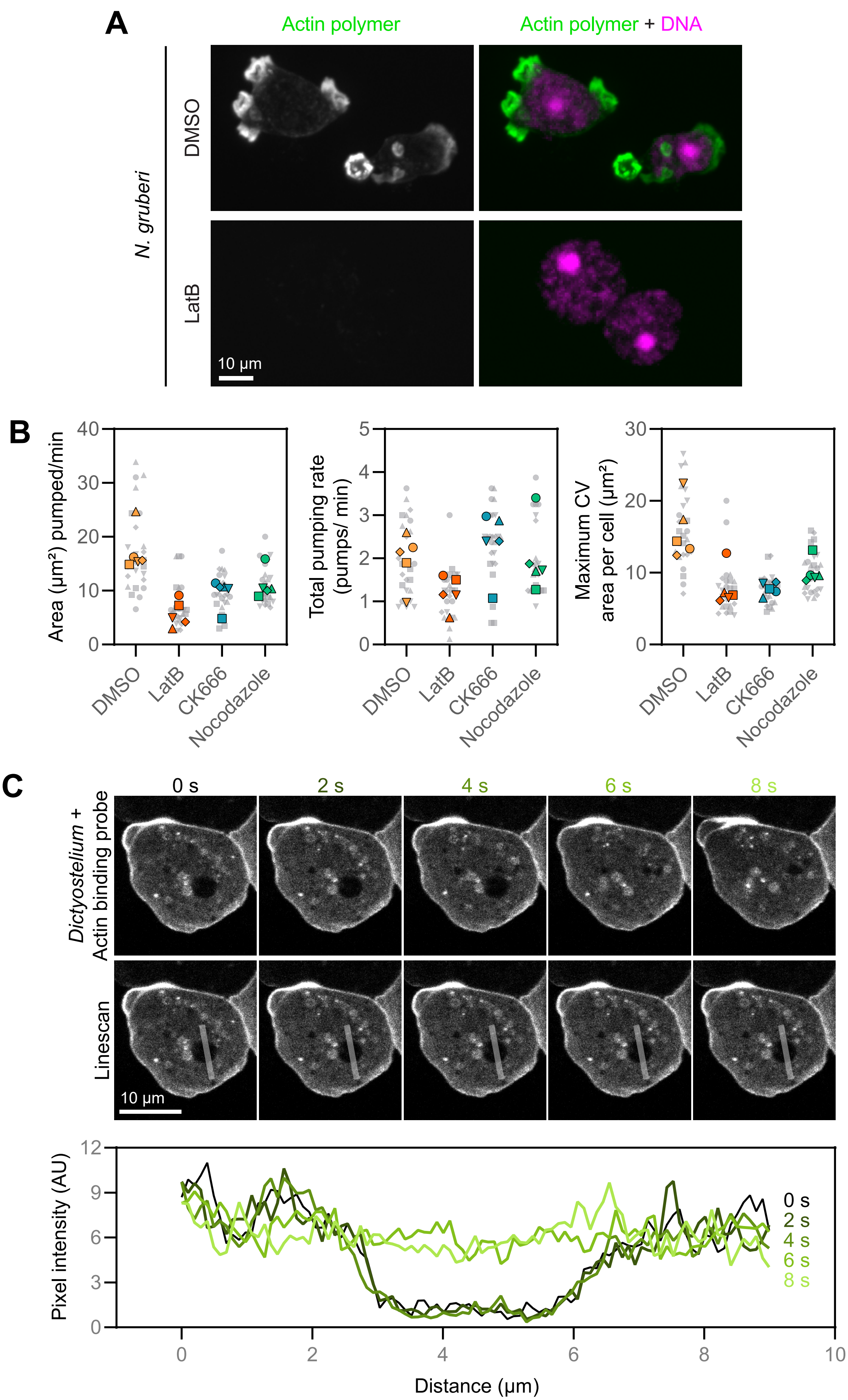

**Figure S3. *Naegleria* and *Dictyostelium* contractile vacuole emptying can occur in the absence of cytoskeletal polymers.** (**A**) *N. gruberi* cells were treated with DMSO or Latrunculin B, then fixed and stained for actin polymer (Alexa-488 conjugated phalloidin) and DNA (DAPI). Maximum intensity projections of representative cells are shown, adjusted to the same brightness and contrast settings. (**B**) The same DMSO and Latrunculin data shown in **Fig. 3C** is shown within the context of a wider drug panel (including the Arp2/3 complex inhibitor CK-666 and the microtubule inhibitor Nocodazole). Graphs show the area pumped per minute by *Dictyostelium* contractile vacuoles, the number of pumps per minute, and the maximum contractile vacuole area per cell. (**C**) A *D. discoideum* cell expressing a fluorescently-labeled actin-binding protein is shown through a pumping cycle. Line scans bisecting a contractile vacuole show no enrichment of actin around the pumping bladder.

**A**

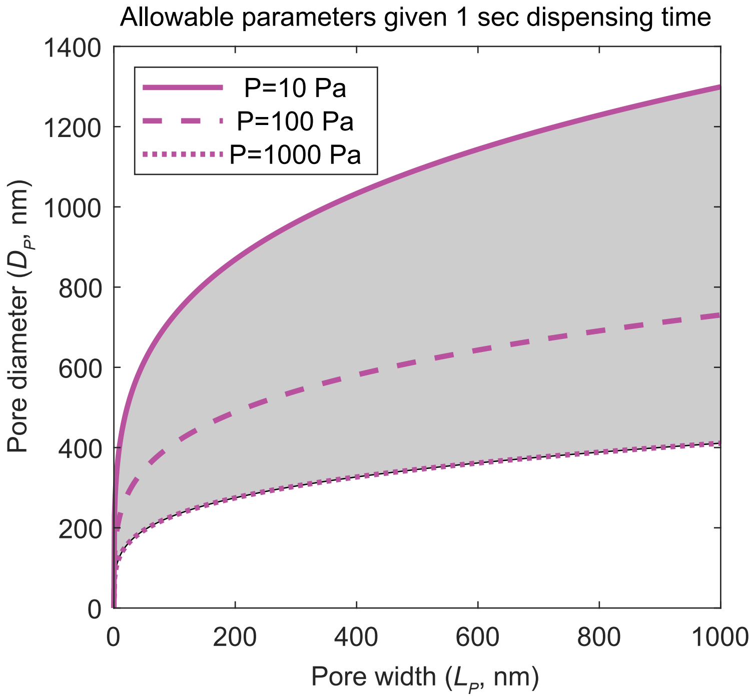

**B**

| Notation | Meaning | Value | Source |
|---|---|---|---|
| $\eta_W$ | Viscosity of water at 25 ºC | 0.9 x $10^{-6}$ pN ms $nm^{-2}$ | -- |
| $H_V$ | Height of the vacuole | 10 μm | This work |
| $D_V$ | Diameter of the vacuole | 10 μm | This work |
| $\Delta P$ | Cytoplasmic pressure | ~10-1000 Pa | Petrie and Koo, Curr. Protoc. Cell Biol., 2014 |
| $L_P$ | Width of the pore | 1 nm - 1 μm | Estimated to be larger than the thickness of the membrane but smaller than the typical z-resolution for light microscopy |
| $D_P$ | Diameter of the pore | ~ 70 nm - 1.3 μm | Derived such that expulsion time = 1 second (See Text S2) |

**Figure S4. Estimating contractile vacuole pore size.** (**A**) Model parameters for the cytoplasmic pressure model that allow the contractile vacuole to dispense its entire volume in 1 s. The required pore diameter is plotted as a function of the choice of pore width, given a cytoplasmic pressure of 10 Pa (solid line), 100 Pa (dashed line), or 1000 Pa (dotted line). Any reasonable choice of cytoplasmic pressure in between 10 and 1000 Pa (gray shading) allows for a wide range of possible pore diameters and pore depths. (**B**) A table shows the allowable parameters[66] representing the results shown in (A).

### Supplemental Text 1: Modeling contractile vacuole filling

#### 1. Introduction

We propose a model in which contractile vacuole dynamics are driven by the competition between the energetic gain associated with lowering osmotic pressure differences across the contractile vacuole membrane, $\Delta\Pi_{CV}$, and the energetic penalty associated with its surface tension, σ, and the cytosolic elastic energy, $G$.

#### 2. Initial filling of the contractile vacuole network

We hypothesize that contractile vacuole helps maintain a constant cytosolic solute concentration for stable cell functions. Given that environmental solute concentrations rarely change on the time scale of contractile vacuole pumping events (~1 min), a constant cytosolic concentration leads to a constant osmotic pressure across the plasma membrane, $\Delta\Pi_{PM}$. To remove excess water in the cytosolic space, contractile vacuoles maintain a solute concentration higher than the cytosolic concentration, and therefore, establish an osmotic pressure across the contractile vacuole membrane, $\Delta\Pi_{cv}$. Water intake leads to a volume increase in the contractile vacuole network prior to its pumping event. Here we derive a simple relation between the two osmolarity differences during the initial filling stage of the contractile vacuole network. From Darcy's law, the instantaneous flow rate across a semipermeable membrane is given by,

$$q = -\frac{k}{\mu}\Pi$$

where $q$ is the instantaneous flow rate per unit area, $k$ is the membrane permeability, μ is the fluid viscosity, and Π is the osmotic pressure across the membrane.

Therefore, the volumetric flow rates across the plasma membrane and the contractile vacuole membrane are given by,

$$Q_{PM} = A_{PM}\frac{k}{\mu}\Delta\Pi_{PM}$$

$$Q_{CV} = A_{CV}\frac{k}{\mu}\Delta\Pi_{CV}$$

where $Q$ is the instantaneous volumetric flow rate, $A$ is the surface area, and the subscripts $PM$ and $CV$ indicate plasma membrane and contractile vacuole membranes, respectively. The membrane permeabilities for the plasma membrane and the contractile vacuole membranes are assumed to be the same, supported by similar membrane staining patterns via FM4-64.

In this analysis, we assume that the cytosolic volume remains constant, agreeing with the experimental observation that the cell area does not significantly change during the contractile vacuole cycle. A constant cytosolic volume requires equal amounts of influx across the plasma membrane and outflux across the contractile vacuole membrane, or mathematically, $Q_{PM} = Q_{CV}$. Rearranging this steady-state condition, we reached the following relation,

$$\Delta\Pi_{CV} = \frac{A_{PM}}{A_{CV}}\Delta\Pi_{PM} \qquad \text{Eq 1}$$

As shown in Eq 1, the osmotic pressure across the contractile vacuole membrane, $\Delta\Pi_{CV}$, is inversely proportional to the total surface area of the contractile vacuole networks, $A_{CV}$. Assuming that the plasma membrane maintains a constant surface area, $A_{PM}$, and a constant osmotic pressure difference, $\Delta\Pi_{PM}$, having a larger contractile vacuole surface area will allow water to flow into the CV at a lower $\Delta\Pi_{CV}$. This could indicate that having a CV network with more surface area is more efficient at establishing and maintaining an osmotic gradient, as it would require less vacuolar type proton pump activity—and therefore less ATP—than a CV network with less surface area.

##### 3. Orders-of-magnitude estimation of the critical size of contractile vacuole network

As the contractile vacuole network fills, the increased membrane tension and cytosolic pressure result in a prominent energetic penalty. The competition between the energetic gain of lower osmotic pressure and the energetic penalty determines a critical size of the contractile vacuole network. Here we derive an orders-of-magnitude estimation of this critical vacuole size as a function of its environmental and intrinsic parameters.

The cell free energies ($F$) pertinent to our analysis are the energies contributed by the osmotic pressure differences ($\Delta\Pi_{PM}$ and $\Delta\Pi_{CV}$) and the tension of the plasma and contractile vacuole membranes ($\sigma_{PM}$ and $\sigma_{CV}$), as well as the cytosolic elastic energy ($GV$). Collectively, the system free energy is expressed as the following equation,

$$F = -\Delta\Pi_{PM}V_{cytosol} - \Delta\Pi_{CV}V_{CV} + \sigma_{PM}(A_{PM} - A_{PM,0}) + \sigma_{CV}(A_{CV} - A_{CV,0}) + GV_{cytosol}$$

where the cytosolic elastic energy is approximated as a linear function of the cytosol volume via an elastic modulus, $G$, and a resting membrane area is assumed for the plasma membrane, $A_{PM,0}$, and the contractile vacuole membrane, $A_{CV,0}$.

In this orders-of-magnitude analysis, we approximated the cell and the contractile vacuole network as spherical objects with radii of $r_1$ and $r_2$, respectively. Another approximation using cylindrical objects yields a similar trend, but a different functional form, between the vacuole critical size and the osmotic pressure. The resting membrane areas, $A_{PM,0}$ and $A_{CV,0}$, are independent of the instantaneous radii, $r_1$ and $r_2$. The first derivative of the system free energy is written as,

$$dF = -4\pi\Delta\Pi_{PM}\left(r_1^2dr_1 - r_2^2dr_2\right) - 4\pi\Delta\Pi_{CV}r_2^2dr_2 + 8\pi\sigma_{PM}r_1dr_1 + 8\pi\sigma_{CV}r_2dr_2 + 4\pi G(\left(r_1^2dr_1 - r_2^2dr_2\right)$$

Following the steady-state cytosolic volume assumption, we recognize that $dV_{cytosol} = 4\pi\left(r_1^2dr_1 - r_2^2dr_2\right) = 0$ and the elastic energy term vanishes, which suggests that the cytosolic elastic energy does not affect the critical contractile vacuole network size prior to pumping. To obtain the critical size at which the contractile vacuole expels, we calculate $r_2^*$ at which $\frac{dF}{dr_2} = 0$, and it has the following form,

$$r_2^* = \frac{2\sigma_{CV}r_1}{\Delta\Pi_{CV}r_1 - 2\sigma_{PM}} \qquad \text{Eq 2}$$

where we have invoked the relation $\frac{dr_1}{dr_2} = \frac{r_2^2}{r_1^2}$ derived from the steady-state cytosolic volume assumption.

As shown in Eq 2, a lower osmotic pressure across contractile vacuole membrane, $\Delta\Pi_{CV}$, corresponds to a larger contractile vacuole network radius, $r_2^*$, qualitatively agreeing with the simple relation derived in Eq 1. To have a more quantitative intuition of the system, we perform an order-of-magnitude estimation by assuming that $O(\sigma_{cv}) = O(\sigma_{PM}) = 1$ mN/m for membrane tension and $O(r_1) = 10$ μm. From Eq 2, we can estimate that for a wide range of solute concentration differences across the contractile vacuole membrane ($0.1 - 10$ mM), the contractile vacuole radius is predicted to be on the order of 1 μm, or $O\left(r_2^*\right) = 1$ μm, consistent with our experimental observations.

To further link this model with experimental data, we derive a relation between $r_2^*$ and $\Delta\Pi_{PM}$. A quadratic equation in $r_2^*$ is derived by combining Eq 1 and Eq 2 as the following,

$$r_2^{\ 2} + r_1 r_2 + \frac{\Delta\Pi_{PM}}{2\sigma} r_1^{\ 3} = 0$$

To simplify the model, here we assume that $\sigma_{PM} = \sigma_{CV}$ as the contractile vacuole and the plasma membrane exhibit similar membrane staining patterns, indicating that these two membranes possess similar compositions, and therefore, similar surface energies.

Taking the positive solution for the quadratic equation, we derived the following expression for $r_2^*$,

$$r_2^{\ *} = \frac{r_1}{2}\left(\sqrt{1 + \frac{2\Delta\Pi_{PM}}{\sigma} r_1} - 1\right) \qquad \text{Eq 3}$$

As shown in Eq 3, a higher environmental solute concentration, corresponding to a lower solute concentration difference and subsequently a lower $\Delta\Pi_{PM}$, will result in a decrease in $r_2^*$, or a smaller contractile vacuole network. This qualitatively agrees with the experimental observation that an increase in the environmental sorbitol concentration results in smaller contractile vacuole networks prior to pumping, suggesting that this simple model can capture the essential dynamical processes in contractile vacuole networks.

### Supplemental Text 2: Modeling contractile vacuole emptying

### 1 Introduction

Here, we develop a physical model for how a cell rapidly expells the contents of its contractile vacuole. This works extends off of previously-published models of contractile vacuole expulsion (Naitoh et al., 1997) and applies them to the specific geometry of cells under confinement. The following section presents the basic assumptions of the model. The subsequent sections explore different possible drivers of vacuole expulsion, and their predictions for experimental data.

### 2 Basic model assumptions

Here we present the basic assumptions of the model.

#### 2.1 The base model

In all versions of the model, we assume that all of the contents of the vacuole exit through a single pore of constant radius and thickness. The fluid in the vacuole is assumed to be an incompressible Newtonian fluid (water) engaging in laminar flow through the pore. The flow through this pore is therefore determined by the Hagen–Poiseuille law for laminar flow through a cylinder:

$$\Delta P = \frac{8\pi\eta L_P v_P}{A_P}, \tag{1}$$

where $\Delta P$ is the pressure difference between either side of the pore, $\eta$ is the dynamic viscosity of the fluid, $L_P$ is the thickness of the pore, $v_P$ is the average velocity of the fluid through the pore, and $A_P$ is the cross-sectional area of the pore. Solving for the velocity of the fluid...

$$v_P = \frac{A_P \Delta P}{8\pi\eta L_P}. \tag{2}$$

The volumetric flow rate through the pore is thus the average fluid velocity mulitplied by the cross-sectional area of the pore, or $\frac{dV}{dt} = Av$. This gives a volumetric flow rate of...

$$\begin{aligned} \frac{dV}{dt} &= A_P v = \frac{A_P^2 \Delta P}{8\pi\eta L_P}. \\ &= \gamma \Delta P \end{aligned} \tag{3}$$

where $\gamma = \frac{A_P^2}{8\pi\eta L_P}$ is the inverse of the resistance to flow through the pore.

To calculate the volume of the vacuole over time, one simply needs to integrate the amount of volume lost over time, and subtract it from the initial volume:

$$\begin{aligned} V(t) &= V_0 - \int_0^t \frac{dV}{dt} \\ &= V_0 - \int_0^t \gamma \Delta P \end{aligned} \tag{4}$$

where $A_P$, $L_P$, and $\eta$ are constants. At this point, where subsequent models can diverge is how $\Delta P$ depends on other model parameters (such as the volume, area, or diameter of the vacuole), which are themselves changing over time as the vacuole expells its contents. Further subsections will explore these different choices for $\Delta P(t)$.

##### 2.2 Validation of laminar flow

Laminar flow is defined as a flow in which fluid particles follow along smooth paths in layers, where each layer moves smoothly past the adjacent layers with little or no mixing. For flow through a cylinder, laminar flow occurs when a dimensionless parameter called the Reynolds number $Re$ becomes smaller than $\approx$ 2,100. The Reynolds number for a cylinder is defined as:

$$Re = \frac{\rho v D}{\eta} \tag{5}$$

where $\rho$ is the density of the fluid, $v$ is the mean speed of the fluid, $D$ is the diameter of the cylinder, and $\eta$ is the dynamic viscosity of the fluid. For the case of the vacuole: The density of water is $\rho_w = 10^3 \frac{kg}{m^3} = 10^{-24} \frac{kg}{nm^3}$, the diameter of the pore is at largest $D = 1\mu m = 10^3 nm$, and the dynamic viscosity of water is temperature-dependent $\eta = 2.414 \cdot 10^{-8} \cdot 10^{\frac{247.8}{(T_K - 140)}}$, which is $9 \cdot 10^{-7} \frac{pN \cdot ms}{nm^2}$ at 25C and $7 \cdot 10^{-7} \frac{pN \cdot ms}{nm^2}$ at 37C. At a velocity of $v = 10 \frac{nm}{ms}$ (exiting the $10\mu m$ vacuole in 1 second), the Reynolds number would be $10^{-5}$, so we are indeed at low Reynold's number.

#### 3 Cytoplasmic pressure model

##### 3.1 Model assumptions

Here, we take the base model presented in Section 2.1, and proceed assuming cytoplasmic pressure drives the contents out of the vacuole. In particular, we assume the pressure inside the vacuole equals the cytoplasmic pressure, and that the cytoplasmic pressure is constant throughout expulsion.

We can assume cytoplasmic pressure is constant because the following logic: (1) The only way for cytoplasmic pressure to change is for the concentration of the cytoplasm to change; (2) The only way for the concentration of the cytoplasm to change is by a flow of water across the cell or vacuole membrane. (3) Because the rate of vacuole expulsion is roughly sixty times faster than the rate of vacuole swelling, we know the flow of water through the open pore is much faster than the rate at which water flows across the intact cell and vacuole membranes; (4) For simplicity, we will thus assume the flow of water across the cell and vacuole membranes is approximately zero during the expulsion phase, and the cytoplasmic pressure remains constant.

##### 3.2 Vacuole volume over time

Mathematically, the simplest possible assumption one can make for $\Delta P$ in Eq. 4 is that it is constant in time. Because $A_P$, $L_P$, and $\eta$ are also constants, the volumetric flow rate is constant. In this case, the total volume in the vacuole over time is...

$$\begin{aligned} V(t) &= V_0 - \int_0^t \gamma \Delta P \\ &= V_0 - \gamma \Delta P t. \end{aligned} \tag{6}$$

#### 3.3 Time to dispense entire volume

To solve for the time is takes to completely empty the vacuole, we can set $V(t_E) = 0$ and solve for $t_E$:

$$\begin{aligned} 0 &= V_0 - \gamma \Delta P t_E \\ t_E &= \frac{V_0}{\gamma \Delta P} \\ &= \frac{V_0 8\pi\eta L_P}{A_P^2 \Delta P} \end{aligned} \tag{7}$$

We can now plug in estimates for a few parameters and get a sense of how large the cytoplasmic pressure has to be in order to expel the contents of a vacuole in $\approx 1$ sec. Let's say the height of the vacuole is $H \approx 10^4 nm$, and the radius is $R = 5 \cdot 10^3 nm$, making the initial volume $V_0 \approx 10^{12} nm^3$. We can also estimated that the diameter of the pore is $D_P \approx 10^3 nm$, the area of the pore is $A_P = \pi((D_P/2)^2 \approx 10^6 nm^2$, and the length of the pore is $L_P \approx 10^3 nm$. Assuming the contents of the vacuole behave as water, the dynamic viscosity of water is temperature-dependent $\eta = 2.414 \cdot 10^{-8} \cdot 10^{\frac{247.8}{(T_K - 140)}}$, which is $\approx 10^{-6} \frac{pN \cdot ms}{nm^2}$ at typical experimental temperatures ($\approx 20C - 40C$). Plugging these values in gives $t = \frac{3*10^{-2} pNms/nm^2}{\Delta P}$. If a typical vacuole expells it's fluid in $\approx 1$ sec, then we can deduce $\Delta P \approx 3 \cdot 10^{-5} pN/nm^2$ (30 Pa, or $3 \cdot 10^{-4}$ atm).

This value is actually at the lower end of measured intracellular pressures ($\approx$ 10 - 1000 Pa, according to `https://doi.org/10.1002%2F0471143030.cb1209s63`), and is well below the typical rupture force of the plasma membrane ($\approx 10kPa$, according to `https://doi.org/10.1016%2Fj.bpj.2016.11.001`). Therefore a reasonable range of cytoplasmic pressures are expected to be sufficient to drive contractile vacuole expulsion.

##### 3.3.1 Useful conversions

$$\begin{aligned} 1Pa &= \frac{N}{m^2} \\ &= \frac{10^{12} pN}{10^{18} nm^2} \\ &= 10^{-6} \frac{pN}{nm^2} \end{aligned} \tag{8}$$

In other words, $\frac{pN}{nm^2} = 1MPa$ (about 10X atmospheric pressure).

#### 3.4 Predictions for 2D microscopy data

Typically in microscopy experiements, vacuole size is tracked by its cross-sectional area as viewed from above, rather than the full 3D volume. Here we derive Eq. 6 in terms of the cross-sectional area, to make predictions for experimental data.

##### 3.4.1 Spherical vacuole

As an exercise, as first assume the vacuole takes the shape of a sphere. Rewriting the volume in terms of its diameter, $V = \frac{4}{3}\pi R^3 = \frac{1}{6}\pi D^3$, the change in the vacuole diameter over time is predicted to be:

$$\begin{aligned} \frac{1}{6}\pi D^3 &= \frac{1}{6}\pi D_0^3 - \gamma \Delta P t \\ D^3 &= D_0^3 - \frac{6\gamma\Delta P}{\pi} t \\ &= D_0^3 - \frac{3A_P^2 \Delta P}{4\pi^2 \eta L_P} t \end{aligned} \tag{9}$$

where the diameter cubed scales linearly with time.

##### 3.4.2 Cylindrical vacuole

In a context where the cell is confined to a defined height in the out-of-plane dimension, the vacuole takes the approximate shape of a cylinder. Rewriting the volume in terms of the cylindrical barrel's diameter, $V = \pi R^2 H = \frac{1}{4}\pi D^2 H$, and the the change in vacuole diameter over time is thus...

$$\begin{aligned}\frac{1}{4}\pi D^2 H &= \frac{1}{4}\pi D_0^2 H - \gamma \Delta P t \\ D^2 &= D_0^2 - \frac{4\gamma\Delta P}{\pi H}t. \\ &= D_0^2 - \frac{A_P^2 \Delta P}{2\pi^2 \eta L_P H}t.\end{aligned} \tag{10}$$

where the diameter squared scales linearly with time. Perhaps more usefully we can re-write this in terms of the cross-sectional area of the cylinder, where $V = AH$...

$$\begin{aligned}AH &= A_0 H - \frac{A_P^2 \Delta P}{8\pi\eta L_P}t \\ A &= A_0 - \frac{A_P^2 \Delta P}{8\pi\eta L_P H}t\end{aligned} \tag{11}$$

Thus if cytoplasmic pressure is driving vacuole expsulsion, we expect to find the cross-sectional area to scale linearly with time.

### 4 Membrane tension model

#### 4.1 Model assumptions

Here, we take the base model presented in Section 2.1, and proceed assuming membrane tension drives the contents out of the vacuole.

#### 4.2 Vacuole volume over time

If the pressure inside the vacuole is dominated by membrane tension, then we can define the pressure in the vacuole using Laplace's law.

##### 4.2.1 Spherical vacuole

For the pressure inside a sphere, Laplace's law is as follows:

$$\Delta P = \frac{4T}{D_{\mathrm{V}}} \tag{12}$$

where $T$ is the membrane tension, which is constant, and $D_V$. Re can start by rewriting the volume $V$ in terms of $D$...

$$\begin{aligned}V &= \frac{4}{3}\pi\left(\frac{D_{\mathrm{V}}}{2}\right)^3 \\ &= \frac{\pi D_{\mathrm{V}}^3}{6} \\ dV &= \frac{\pi D_{\mathrm{V}}^2}{2}dD\end{aligned} \tag{13}$$

And then rewriting $\frac{dV}{dt}$ in terms of $D$.

$$\begin{aligned}\frac{dV}{dt} &= -\gamma\Delta P \\ \frac{\pi D_{\mathrm{V}}^2}{2}\frac{dD}{dt} &= -\gamma\frac{4T}{D_{\mathrm{V}}}\end{aligned} \tag{14}$$

Isolating $D_V$ and $t$ on either side

$$D_{\mathrm{V}}^3 dD = -\frac{8\gamma T}{\pi} \tag{15}$$

and integrating...

$$\int_{D_0}^{D} D_{\mathrm{V}}^3 dD = -\int_0^t \frac{8\gamma T}{\pi}$$
$$\frac{D^4}{4} - \frac{D_0^4}{4} = -\frac{8\gamma T}{\pi} t \tag{16}$$
$$D^4 = D_0^4 - \frac{32\gamma T}{\pi} t$$

###### 4.2.2 Cylindrical vacuole

For the pressure inside a cylinder, Laplace's law is as follows:

$$\Delta P = \frac{2T}{D_{\mathrm{V}}} \tag{17}$$

where the pressure inside a cylinder differs only by a factor of 2 from that of a sphere of equivalent diameter. Following the same logic as for a sphere, we find...

$$D^4 = D_0^4 - \frac{16\gamma T}{\pi} t \tag{18}$$

Thus if membrane tension is driving vacuole expsulsion, we expect to find the (cross-sectional area)$^2$ to scale linearly with time.
